# Onco-Circuit Addiction and Onco-Nutrient mTORC1 Signaling Vulnerability in a Model of Aggressive T Cell Malignancy

**DOI:** 10.1101/2024.04.03.587917

**Authors:** Xinxin Wang, Andrew E. Cornish, Mytrang H. Do, Julia S. Brunner, Ting-Wei Hsu, Zijian Xu, Isha Malik, Chaucie Edwards, Kristelle J. Capistrano, Xian Zhang, Mark H. Ginsberg, Lydia W.S. Finley, Megan S. Lim, Steven M. Horwitz, Ming O. Li

## Abstract

How genetic lesions drive cell transformation and whether they can be circumvented without compromising function of non-transformed cells are enduring questions in oncology. Here we show that in mature T cells—in which physiologic clonal proliferation is a cardinal feature— constitutive *MYC* transcription and *Tsc1* loss in mice modeled aggressive human malignancy by reinforcing each other’s oncogenic programs. This cooperation was supported by MYC-induced large neutral amino acid transporter chaperone SLC3A2 and dietary leucine, which in synergy with *Tsc1* deletion overstimulated mTORC1 to promote mitochondrial fitness and MYC protein overexpression in a positive feedback circuit. A low leucine diet was therapeutic even in late-stage disease but did not hinder T cell immunity to infectious challenge, nor impede T cell transformation driven by constitutive nutrient mTORC1 signaling via *Depdc5* loss. Thus, mTORC1 signaling hypersensitivity to leucine as an onco-nutrient enables an onco-circuit, decoupling pathologic from physiologic utilization of nutrient acquisition pathways.

## INTRODUCTION

Oncogenesis is a multistep process.^1^ Among the many reasons for this, the most well-described is a requirement for multiple genetic lesions to co-occur in an individual cell.^2^ Such oncogenic events must cooperate to induce cell transformation, either synergistically to hyperactivate growth and proliferation programs or consecutively to disable cellular breaks activated in response to earlier oncogenic lesions. Cancer cell dependence on a particular oncogenic lesion for maintenance, and not just initiation, of the malignant phenotype is referred to as “oncogene addiction” and has provided a conceptual foundation for the considerable clinical success of targeted therapies.^3^ Along with cooperative genetic events, cancer cells also adapt cellular pathways that lack mutations. Their heightened dependence on such non-mutated pathways is referred to as “non-oncogene addiction” and has provided a conceptual foundation for therapeutic strategies including some types of synergistic lethality.^4^

How individual oncogenic genetic lesions, or ‘onco-genotype’, act in concert with one another as well as non-mutated pathways to achieve an oncogenic functional program, or ‘onco-phenotype’, varies considerably. Comparatively straightforward examples include gain-of-function mutations in kinases such as *PI3KCA* leading directly to aberrant signaling,^5^ or loss-of-function mutations in tumor suppressors such as *TP53* leading to DNA instability or evasion of cell cycle arrest and apoptosis.^6^ A somewhat more convoluted example is that of the transcription factor MYC, which is dysregulated in up to 70% of human cancers with constitutive *MYC* transcription frequently driven by genomic amplification and chromosomal translocation.^7,8^ However, translational and post-translational mechanisms further control levels of MYC protein, which binds approximately 15% of the genome to orchestrate transcriptional programs essential for cell transformation.^9,10^ Elucidating how high MYC protein levels are induced, and how MYC-driven transcriptional programs collaborate with other cellular processes to maintain a cancerous cell state, are of particular importance given the noted difficulty in therapeutically targeting MYC itself.^11^

Likewise, many oncogenic pathways intersect with the mammalian/mechanistic target of rapamycin complex 1 (mTORC1) in an intricate manner. mTORC1 regulates macromolecule biosynthesis and cell growth by assimilating extracellular and intracellular information, in particular growth factor signaling and nutrient availability via the Rheb and Rag GTPases, respectively.^12^ Many of the most frequently activated oncogenic signaling pathways in human cancer, including PI3K/AKT and RAS-RAF-MEK-ERK, converge to inactivate the TSC complex, the GTPase activating protein (GAP) for Rheb, and simulate persistent growth factor signaling in a cell-autonomous fashion.^13^ Yet, translocation of mTORC1 to the lysosomal surface for activation by Rheb requires adequate nutrients, particularly amino acids, sensed in part through the GATOR1 complex, the GAP for RagA/B GTPase.^14^ Elucidating how mTORC1 signaling is induced, and how it collaborates with other oncogenic lesions to drive cell transformation, are of great importance given limited use of mTORC1 inhibitors as cancer monotherapies due to drug toxicity and resistance.^15^

In this study, we show that constitutive *MYC* transcription and deletion of the TSC complex component gene *Tsc1* cooperated in mice to induce T cell transformation and modeled aggressive human disease. MYC elevated amino acid transporter expression to promote amino acid uptake and nutrient mTORC1 signaling, which acted in concert with *Tsc1* loss to induce mTORC1 hyperactivation. mTORC1 reciprocally facilitated stable accumulation of MYC protein, together forming a positive feedback circuit culminating in aggressive malignancy. The dependence of this circuit on mobilization of amino acid membrane transporters was illustrated by ablation of the large neutral amino acid transporter chaperone SLC3A2, which impeded transformation by compromising the mitochondrial fitness of inchoate cancer cells. This phenotype was also achieved with a low leucine diet, which improved survival even when initiated after the development of advanced disease. Leucine deprivation did not impair the T cell response to pathogen challenge, nor did it affect T lymphoma development caused by activation of the nutrient sensing branch of mTORC1 via loss of the GATOR1 component gene *Depdc5*. Together, these findings reveal that nutrient mTORC1 signaling serves a rate-limiting role for oncogenic leucine acquisition via SLC3A2 in the *MYC/Tsc1* model, and targeting such ‘onco-circuit’ and ‘onco-nutrient’ addiction heralds a new conceptual framework for cancer therapy.

## RESULTS

### Constitutive transcription of *MYC* or a T58A *MYC* mutant does not readily transform T cells

To define how oncogenic pathways cooperate to induce cell transformation, we set forth to study T cell malignancies. Clonal expansion is an essential aspect of normal T cell physiology, thus studying T cell oncogenesis may offer insights into therapeutically exploitable molecular programs that distinguish the clonal propagation of cancer cells from that of non-transformed healthy cells.

In addition, despite being a rare and heterogeneous cancer type accounting for ∼10-20% of non-Hodgkin lymphomas worldwide, peripheral T cell lymphomas (PTCLs) are highly aggressive and frequently refractory to treatment, and patients with PTCL have not benefited from new therapeutic modalities such as targeted therapy and immunotherapy that have had success in other cancer indications.^16^

Genomic profiling studies have revealed that genes encoding components of growth factor signaling pathways downstream of T cell receptor signaling, especially those associated with the PI3K/AKT signaling axis, are frequently mutated in human PTCLs.^17,18^ Furthermore, the *MYC* oncogene is amplified or overexpressed in a large fraction of PTCL tumors in association with poor patient survival.^18–20^ To model PTCL in mice, the murine ortholog of recurrently mutated human tumor suppressor gene *PTEN* encoding phosphatidylinositol-3,4,5-triphosphate 3-phosphatase, a pivotal PI3K signaling inhibitor, was conditionally ablated at CD4^-^CD8^-^ or CD4^+^CD8^+^ stages of T cell development by LckCre or CD4Cre, respectively, which led to fulminant T cell transformation and mouse lethality within 10-16 weeks of age.^21,22^ Intriguingly, close examination of this CD4Cre model showed that all lymphoma T cells harbored t(14:15) chromosomal translocations that drive *Myc* transcription from the *Tcra* gene locus.^23^ Importantly, deletion of *Myc* corrected the lymphoma phenotype,^24^ demonstrating an essential role for the translocated *Myc* allele in T cell lymphomagenesis originated from PTEN loss.

To investigate whether PTEN loss induced T cell transformation principally through promoting constitutive *MYC* transcription, we crossed CD4Cre mice with *Rosa26^MYC/+^*mice harboring a *CAG* promoter-driven human *MYC* allele preceded by a *LoxP* site-flanked translation stop sequence and followed by a signaling-deficient human *CD2* reporter in the constitutively active *Rosa26* locus.^25^ This allele temporally recapitulates MYC induction in the PTEN model more closely than previously reported transgenic mice with sustained expression of MYC in T cells.^26,27^ Of note, unlike the PTEN model, CD4Cre*Rosa26^MYC/+^* mice did not manifest a fulminant lethality phenotype (Figure S1A), suggesting that signaling event(s) downstream of PTEN loss may synergize with constitutive *MYC* transcription to induce T cell transformation.

One such candidate event is that PTEN loss-induced PI3K/AKT signaling may stabilize MYC protein by preventing GSK3B-induced Thr58 phosphorylation, which triggers MYC ubiquitination and proteasome-mediated degradation.^28,29^ To explore this aspect of MYC regulation in lymphoma development, we crossed CD4Cre mice with *Igs2^MYC(T58A)/+^*mice carrying a *CAG* promoter-driven *MYC* Thr58 to Ala mutant allele preceded by a *LoxP* site-flanked translation stop sequence in the constitutively active *Igs2* locus.^30^ Unexpectedly, CD4Cre*Igs2^MYC(T58A)/+^*mice similarly did not succumb to early lethality (Figure S1A), implying that other or additional signaling event(s) triggered by PTEN deficiency may cooperate with constitutive *MYC* transcription to induce T cell malignancy.

### *MYC* transcription synergizes with *Tsc1* loss to drive a T cell lymphoproliferative disease

In addition to repressing GSK3B, PTEN loss-induced PI3K/AKT signaling may phosphorylate and inactivate the TSC complex that functions as a GAP for the mTORC1 activator small GTPase Rheb.^31^ Of note, aside from *PTEN* loss and other PI3K/AKT-activating mutations, recurrent loss-of-function mutations of the TSC complex itself have been detected in human PTCLs.^18^ To investigate whether TSC inactivation might cooperate with constitutive *MYC* transcription to induce T cell transformation, we crossed CD4Cre*Rosa26^MYC/+^* mice with *Tsc1^fl/fl^* mice to block TSC signaling in the T cell lineage. While T cell-specific ablation of Tsc1 alone did not affect mouse survival, CD4Cre*Rosa26^MYC/+^Tsc1^fl/fl^* mice were visibly distressed and succumbed to death by 8 weeks of age (Figure 1A). Histopathology and flow cytometry analyses revealed extensive T cell expansion and/or infiltration in lymphoid and non-lymphoid organs including spleen, lymph nodes, and liver as well as in blood in CD4Cre*Rosa26^MYC/+^Tsc1^fl/fl^*, but not control, mice (Figure S1B, S1C). Similar to T cell lymphomas in the PTEN model^23^ as well as the majority of human T cell lymphoma cases,^16^ the expanded T cells from CD4Cre*Rosa26^MYC/+^Tsc1^fl/fl^* mice had a predominant CD4^+^CD8^-^ phenotype (Figure S1D). Conversely, CD4 and CD8 T cell profiles were largely unperturbed in CD4Cre*Rosa26^MYC/+^*mice, and CD4Cre*Tsc1^fl/fl^* mice had decreased CD8^+^ T cells as previously reported^32^ (Figure S1D). The aberrantly expanded CD4^+^ T cells were positive for the cell proliferation marker Ki-67 but did not express the regulatory T (Treg) cell marker Foxp3 (Figure S1E). These findings reveal strong synergy between constitutive *MYC* transcription and *Tsc1* loss in driving a lethal T cell lymphoproliferative disease and suggest that PTEN deficiency largely acts through the PI3K/AKT/TSC signaling axis to cooperate with MYC and induce T cell transformation.

**Figure 1.**
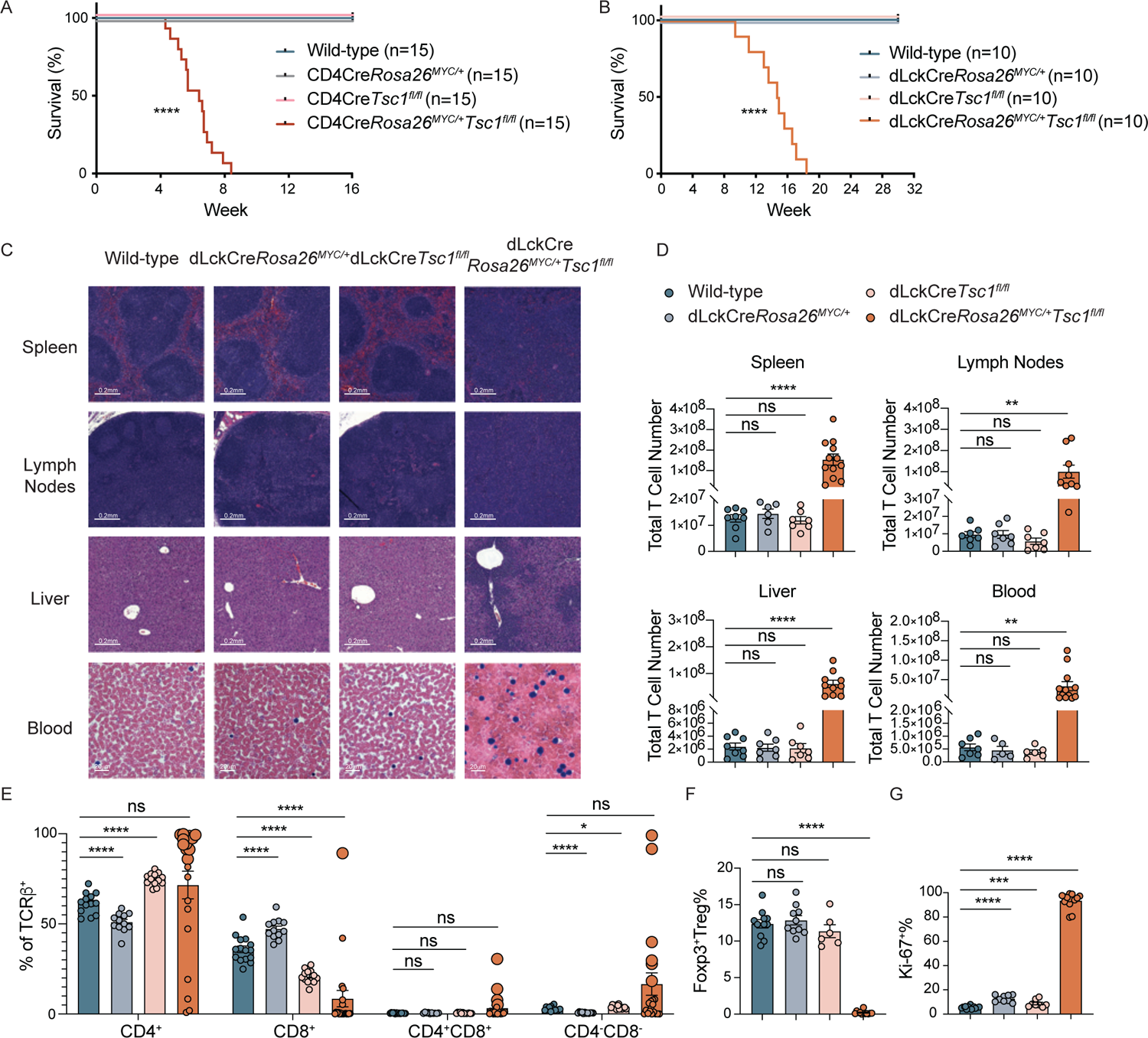
Constitutive *MYC* transcription synergizes with *Tsc1* deletion in T cells to result in lethality associated with a lymphoproliferative T cell phenotype. (A) Survival curves for littermate wild-type control, CD4Cre*Rosa26^MYC/+^*, CD4Cre*Tsc1^fl/fl^*, and CD4Cre*Rosa26^MYC/+^Tsc1^fl/fl^* mice. (B) Survival curves for littermate wild-type control, dLckCre*Rosa26^MYC/+^*, dLckCre*Tsc1^fl/fl^*, and dLckCre*Rosa26^MYC/+^Tsc1^fl/fl^* mice. (C) Rows 1-3: Representative images of hematoxylin and eosin (H&E) staining of spleen, lymph node, and liver in wild-type, dLckCre*Rosa26^MYC/+^*, dLckCre*Tsc1^fl/fl^*, and dLckCre*Rosa26^MYC/+^Tsc1^fl/fl^* mice. Original magnification, 21x. Scale bar 0.2 mm. Row 4: Representative images of Giemsa stain of blood from wild-type, dLckCre*Rosa26^MYC/+^*, dLckCre*Tsc1^fl/fl^*and dLckCre*Rosa26^MYC/+^Tsc1^fl/fl^* mice. Original magnification, 63x. Scale bar 20 μm. (D) Quantification of total T cell numbers in spleen, lymph nodes, liver and blood from wild-type, dLckCre*Rosa26^MYC/+^*, dLckCre*Tsc1^fl/fl^*, and dLckCre*Rosa26^MYC/+^Tsc1^fl/fl^*mice. (E) Frequencies by flow cytometry of CD4^+^, CD8^+^, CD4^+^CD8^+^, and CD4^-^CD8^-^ T cells in spleen from wild-type, dLckCre*Rosa26^MYC/+^*, dLckCre*Tsc1^fl/fl^*, and dLckCre*Rosa26^MYC/+^Tsc1^fl/fl^*mice. Enlarged dots represent statistics of expanded T cell populations. (F) Frequency of Foxp3^+^ Treg cells among CD4^+^ T cells in spleen from wild-type, dLckCre*Rosa26^MYC/+^*, dLckCre*Tsc1^fl/fl^*, and dLckCre*Rosa26^MYC/+^Tsc1^fl/fl^*mice. (G) Frequency of Ki-67^+^ cells among CD4^+^ T cells in spleen from wild-type, dLckCre*Rosa26^MYC/+^*, dLckCre*Tsc1^fl/fl^*, and dLckCre*Rosa26^MYC/+^Tsc1^fl/fl^*mice. All bar graphs are shown as mean ± SEM, each dot representing one mouse. Data are pooled from multiple experiments. [A, B: Log-rank (Mantel Cox) test; D, E, F, G: one-way ANOVA with Tukey’s multiple comparisons test, * = p<0.05, ** = p<0.01, *** = p<0.001, **** = p<0.0001, “ns” = not significant.] See also Figure S1.

While PTCLs are derived from mature post-thymic T lymphocytes,^16^ CD4Cre is expressed at the immature CD4^+^CD8^+^ double positive (DP) stage of T-thymocyte development and may confound the use of CD4Cre*Rosa26^MYC/+^Tsc1^fl/fl^* mice to model human disease. Indeed, although most CD4Cre*Rosa26^MYC/+^Tsc1^fl/fl^* mice displayed thymic involution— likely a consequence of systemic stress—some mice displayed thymic enlargement along with disrupted T cell development (Figure S1F). To address this concern, we used dLckCre mice in which the Cre recombinase is expressed in mature CD4^+^ and CD8^+^ T cells under the control of the distal *Lck* promoter.^33^ As expected, T cell development in dLckCre*Rosa26^MYC/+^Tsc1^fl/fl^*mice was unperturbed (Figure S1G). As in the CD4Cre model, all dLckCre*Rosa26^MYC/+^Tsc1^fl/fl^*mice succumbed to premature death, though with comparatively delayed kinetics (Figure 1B). This lethal phenotype likewise tracked with expansion and infiltration of highly proliferative CD4^+^Foxp3^-^ T cells in spleen, lymph nodes, liver, and blood (Figure 1C-1G). Thus, constitutive *MYC* transcription cooperates with *Tsc1* deletion in mature T cells to trigger a lethal T cell lymphoproliferative disease.

### Mutually-reinforced MYC induction and mTORC1 activation causes aggressive PTCL

To better characterize this lymphoproliferative disease, we performed bulk RNA-sequencing experiments on hCD2^+^CD4^+^CD25^-^ T cells isolated from end-stage CD4Cre*Rosa26^MYC/+^Tsc1^fl/fl^*and dLckCre*Rosa26^MYC/+^Tsc1^fl/fl^* mice with hCD2^+^CD4^+^CD25^-^ T cells from CD4Cre*Rosa26^MYC/+^* mice as well as CD4^+^CD25^-^ T cells from CD4Cre*Tsc1^fl/fl^* and wild-type mice as controls. Calculation of TCR repertoire clonality using transcripts mapped to the TCR-β CDR3 region^34^ revealed that T cells from CD4Cre*Rosa26^MYC/+^Tsc1^fl/fl^* and dLckCre*Rosa26^MYC/+^Tsc1^fl/fl^* mice were oligoclonal and monoclonal, respectively, while T cells from control mice were polyclonal (Figure 2A, S2A). Principal component analysis (PCA) of transcriptomic data further showed that T cells from CD4Cre*Rosa26^MYC/+^Tsc1^fl/fl^*and dLckCre*Rosa26^MYC/+^Tsc1^fl/fl^* mice were mostly distinct from T cells from control mice (Figure S2B). Of note, pathway analyses with HALLMARK and REACTOME signatures demonstrated downregulation of inflammatory response and immune receptor signaling genes in CD4Cre*Rosa26^MYC/+^Tsc1^fl/fl^*and dLckCre*Rosa26^MYC/+^Tsc1^fl/fl^* T cells (Figure 2B, S2C; Table S1). Indeed, these T cells were largely defective in producing T helper cell cytokines TNFα, IFNγ, IL-17A, or IL-4 compared to T cells from control mice (Figure S2D and data not shown). This clonal expansion accompanied by a deficit of immune effector molecule production suggests that the lethal lymphoproliferative disease is of a malignant rather than autoimmune nature, representing a bona fide peripheral T cell lymphoma.

**Figure 2.**
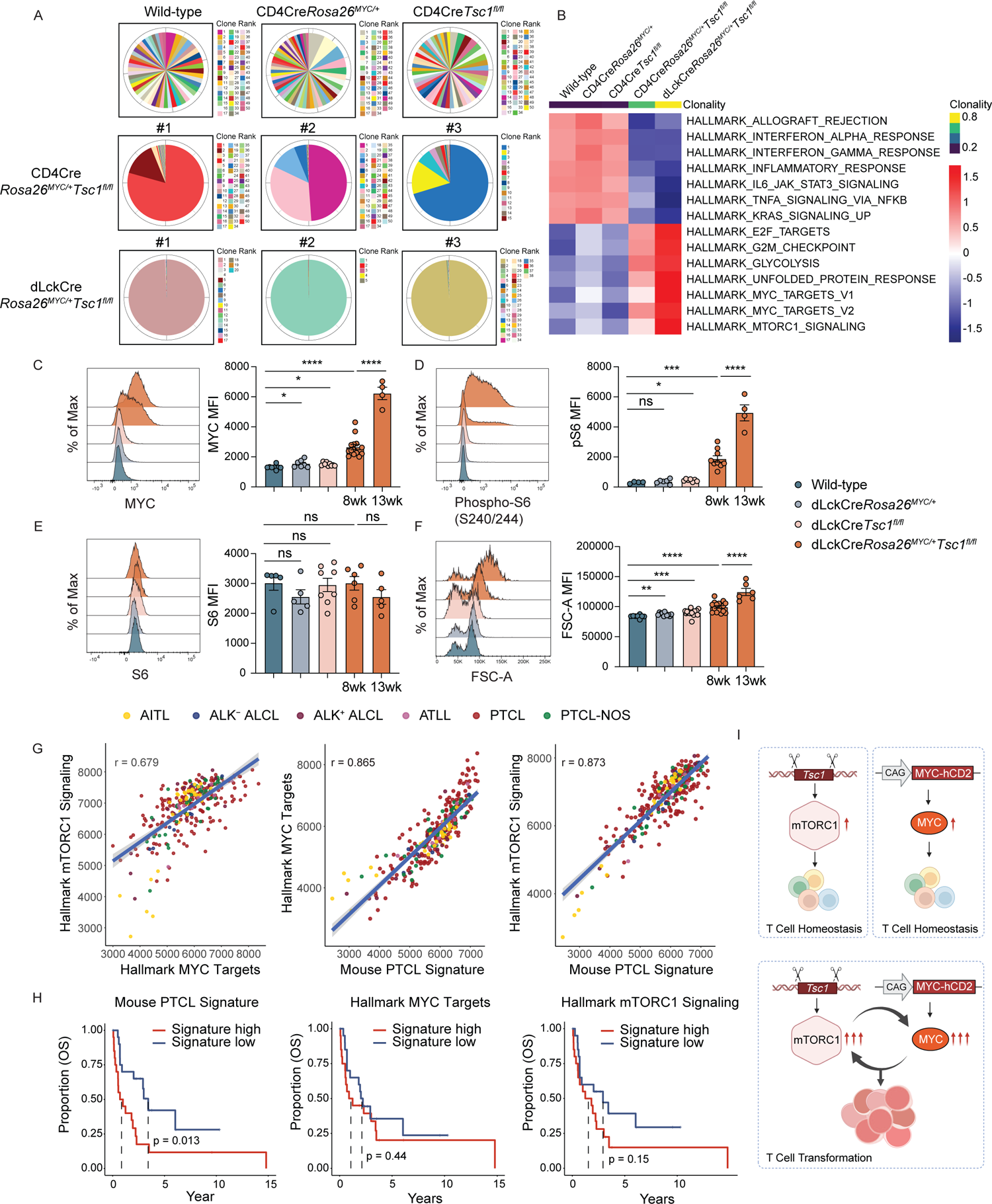
Constitutive *MYC* transcription and *Tsc1* deletion triggers PTCL with its gene expression signature marked patients with poor survival regardless of disease subtypes. (A) Pie charts showing the relative abundance of T cell clones in splenic CD4^+^ T cells isolated from wild-type and CD4Cre*Tsc1^fl/fl^* mice and hCD2^+^CD4^+^ T cells from CD4Cre*Rosa26^MYC/+^*, CD4Cre*Rosa26^MYC/+^Tsc1^fl/fl^*, and dLckCre*Rosa26^MYC/+^Tsc1^fl/fl^*mice. T cell clone is inferred from transcripts mapping to the TCRβ CDR3 region in bulk RNA-sequencing. Plots shown are representative of analysis from two mice each for wild-type, CD4Cre*Rosa26^MYC/+^*, and CD4Cre*Tsc1^fl/fl^*genotypes. Full analysis from three CD4Cre*Rosa26^MYC/+^Tsc1^fl/fl^*and three dLckCre*Rosa26^MYC/+^Tsc1^fl/fl^* mice are also shown. For mice with over 50 unique TCRý CDR3 clones the top 50 are shown, while all clones are shown for mice with under 50 unique TCRβ CDR3 clones. (B) Heatmap of z-scored ssGSEA scores (averaged across 2-3 mice per condition) for select Hallmark signature gene sets. Gene sets included are those differentially enriched between wild-type and dLckCre*Rosa26^MYC/+^Tsc1^fl/fl^*mice, but for heatmap visualization CD4Cre*Rosa26^MYC/+^*, CD4Cre*Tsc1^fl/fl^*, and CD4Cre*Rosa26^MYC/+^Tsc1^fl/fl^*mice are also shown. Genotypes are ordered horizontally by average CDR3 clonality, and gene sets are manually ordered vertically by functional annotation. (C-F) Representative flow cytometric analysis of MYC (C), phosphorylated S6 (D), and S6 (E) protein as well as FSC-A (F) staining (left panels) and MFI Quantification (right panels) in splenic CD4^+^ T cells from wild-type and dLckCre*Tsc1^fl/fl^* mice and hCD2^+^CD4^+^ T cells from dLckCre*Rosa26^MYC/+^* and dLckCre*Rosa26^MYC/+^Tsc1^fl/fl^*mice at 8 weeks of age and dLckCre*Rosa26^MYC/+^Tsc1^fl/fl^*mice at 13 weeks of age. (G) Scatter plot of ssGSEA scores in published PTCL patient microarray dataset for MYC Targets (V2) and mTORC1 Signaling Hallmark gene sets (left); MYC Targets (V2) and Mouse PTCL Signature (middle); and mTORC1 Signaling Signature and Mouse PTCL Signature (right). The Mouse PTCL Signature comprises the human orthologs of differentially expressed genes upregulated in CD4^+^ T cells from dLckCre*Rosa26^MYC/+^Tsc1^fl/fl^* as opposed to wild-type mice. A linear fit and Pearson’s correlation are shown for each scatter plot, and points are colored by the reported PTCL subtype. (H) Kaplan-Meier curve showing overall survival of PTCL-NOS patients grouped by expression (ssGSEA score) above or below the median for the Mouse PTCL Signature (left), Hallmark MYC Targets V2 (middle), and mTORC1 Signaling (right). (I) Schematic illustrating how constitutive *MYC* transcription synergizes with *Tsc1* deletion in a positive feedback circuit to drive peripheral T cell transformation. All bar graphs are shown as mean ± SEM, each dot representing one mouse. Data are pooled from multiple experiments. [C, D, E, F: one-way ANOVA with Tukey’s multiple comparisons test; G: Pearson correlation based on Kolmogorov-Smirnov test; H: Log-rank (Mantel Cox) test, * = p<0.05, ** = p<0.01, *** = p<0.001, **** = p<0.0001, “ns” = not significant.] See also Figure S2.

As expected from this malignant classification, the transcriptome of T cells from CD4Cre*Rosa26^MYC/+^Tsc1^fl/fl^* and dLckCre*Rosa26^MYC/+^Tsc1^fl/fl^* mice was characterized by upregulation of cell cycle-related pathways including E2F targets and G2M checkpoint genes as well as metabolic and stress signaling pathways (Figure 2B, S2C). Unexpectedly, MYC targets and mTORC1 signaling pathway genes were much more strongly induced in CD4Cre*Rosa26^MYC/+^Tsc1^fl/fl^* and dLckCre*Rosa26^MYC/+^Tsc1^fl/fl^* T cells than T cells from CD4Cre*Rosa26^MYC/+^* and CD4Cre*Tsc1^fl/fl^* mice, respectively (Figure 2B). In agreement with these observations, both MYC protein expression and mTORC1 signaling marked by phospho-S6 over total S6 staining were synergistically induced in dLckCre*Rosa26^MYC/+^Tsc1^fl/fl^*T cells relative to single-mutant T cells (Figure 2C-2E). Furthermore, induction of MYC protein expression and activation of mTORC1 signaling co-reinforced each other in a time-dependent manner and tracked with growth of lymphoma T cells (Figure 2C-2F). These findings demonstrate a positive feedback circuit characterized by reinforcement of MYC protein expression and mTORC1 activation in association with aggressive PTCL development in dLckCre*Rosa26^MYC/+^Tsc1^fl/fl^* mice. Using differentially expressed genes that were upregulated in dLckCre*Rosa26^MYC/+^Tsc1^fl/fl^* T cells relative to those in wild-type T cells, we established a mouse PTCL gene signature (Table S1). Analyzing the expression of this signature in published human PTCL microarray data comprising a total of 372 samples,^20^ we found that our mouse PTCL signature correlated highly with the expression of hallmark gene sets for both MYC targets and mTORC1 signaling in PTCL patient samples (Figure 2G). Expression of MYC targets also correlated with mTORC1 signaling genes in human PTCL samples, but to a lesser extent than did either with the mouse PTCL signature, suggesting this signature identified a subgroup of human PTCLs irrespective of disease subtype (Figure 2G). To determine how these signatures tracked with disease severity, human PTCL-NOS samples, the most aggressive subtype, were divided into signature-high and -low groups based on the mouse PTCL signature, MYC targets, or mTORC1 signaling genes, respectively, and survival analyses were performed. There was a trend towards decreased overall survival in both MYC-high versus low and mTORC1 signaling-high versus low groups, though neither met significance (Figure 2H). By contrast, the PTCL signature-high group displayed significantly inferior overall survival compared to the PTCL signature-low group (Figure 2H). Overall, these data suggest that the murine PTCL model, characterized by a positive feedback circuit between induction of MYC protein expression and activation of mTORC1 signaling (Figure 2I), does not model any one subtype of PTCL. Instead, it captures a central molecular program active in the most aggressive cases of PTCL.

### SLC3A2 reinforces the MYC and mTORC1 circuit and PTCL survival in the *MYC/Tsc1* model

We wished to define the molecular architecture of this MYC and mTORC1 circuit operative in dLckCre*Rosa26^MYC/+^Tsc1^fl/fl^*mice and explore its function in PTCL development. As a metabolic regulator, mTORC1 promotes MYC expression by enhancing S6K1-mediated translation initiation,^35^ but how MYC promotes mTORC1 signaling is less well understood. In addition to growth factor signaling that acts through the PI3K/AKT/TSC pathway to control mTORC1 activation via the lysosome-localized Rheb, nutrient availability, particularly of amino acids, regulates mTORC1 lysosomal recruitment and preconditions Rheb-mediated mTORC1 activation.^36,37^ Of note, among the upregulated transcripts in dLckCre*Rosa26^MYC/+^Tsc1^fl/fl^* PTCL T cells encoding plasma membrane proteins were several of the solute carrier (SLC) family transporters including the MYC target *Slc7a5* and *Slc3a2*^38,39^ (Figure 3A, Table S1). The encoded SLC7A5/SLC3A2 (CD98) heterodimer forms the L-type amino acid transporter 1 (LAT1) that promotes cellular uptake of large neutral amino acids with important functions in mTORC1 regulation.^40^ In agreement with the transcriptomic data, CD98 protein was induced in a time-dependent manner in dLckCre*Rosa26^MYC/+^Tsc1^fl/fl^*T cells, kinetically tracking expression of MYC protein (Figure 2C, 3B).

**Figure 3.**
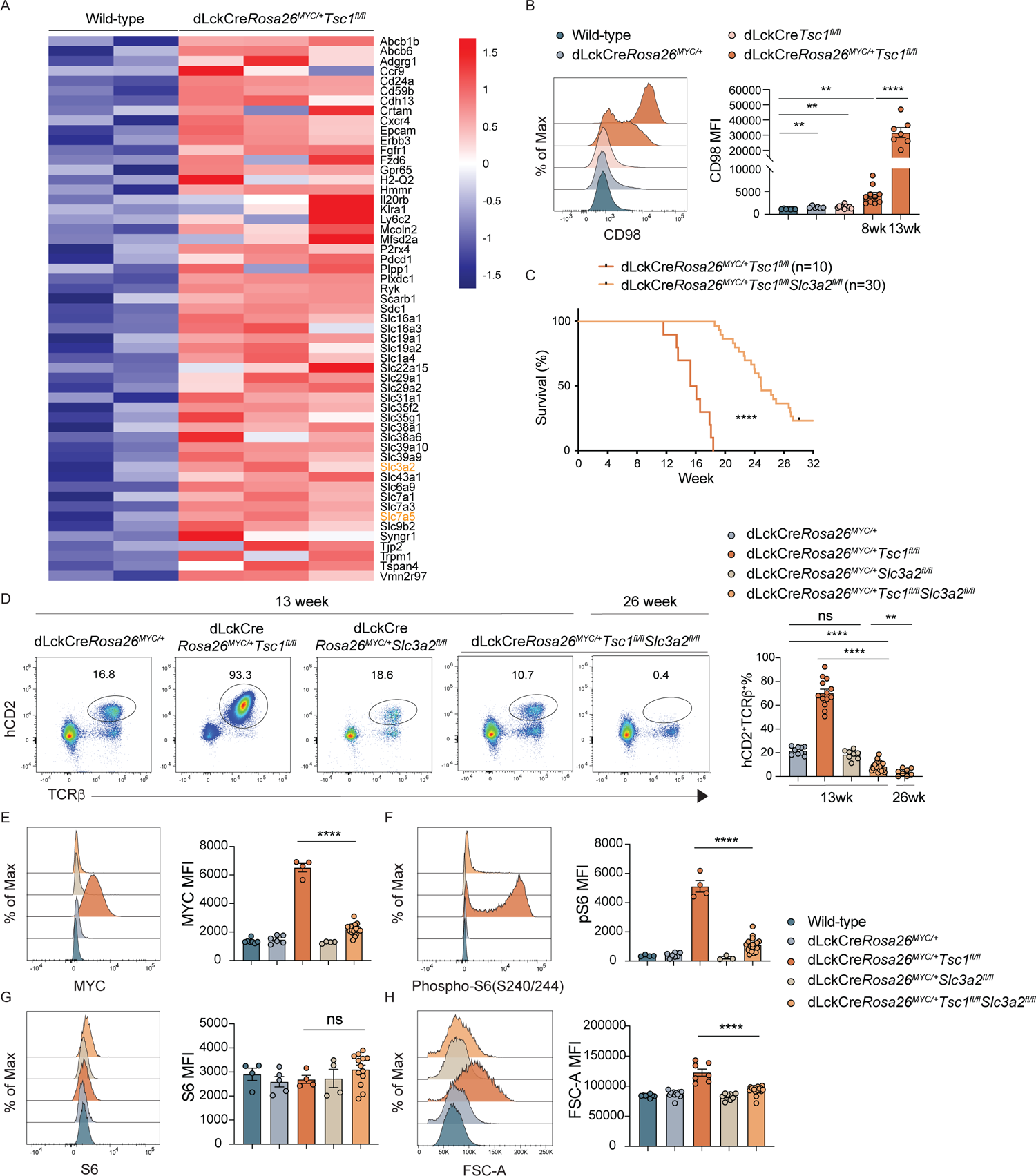
Large neutral amino acid transporter chaperone SLC3A2 is required to sustain high levels of MYC protein expression and mTORC1 signaling for T cell transformation. (A) Heatmap of z-scored expression of cell surface proteins positively differentially expressed in hCD2^+^CD4^+^ T cells from dLckCre*Rosa26^MYC/+^Tsc1^fl/fl^* mice as compared with CD4^+^ T cells from wild-type mice, ordered alphabetically. The two chains of the CD98 heterodimer encoded by *Slc3a2* and *Slc7a5* are highlighted in yellow. (B) Representative flow cytometric analysis (left panel) and MFI quantification (right panel) of CD98 protein expression in CD4^+^ T cells from wild-type and dLckCre*Tsc1^fl/fl^* mice and hCD2^+^CD4^+^ T cells from dLckCre*Rosa26^MYC/+^* and dLckCre*Rosa26^MYC/+^Tsc1^fl/fl^* mice at 8 weeks of age as well as dLckCre*Rosa26^MYC/+^Tsc1^fl/fl^*mice at 13 weeks of age. (C) Survival curves for dLckCre*Rosa26^MYC/+^Tsc1^fl/fl^Slc3a2^fl/fl^*mice compared to dLckCre*Rosa26^MYC/+^Tsc1^fl/fl^* mice. (D) Left panel: Representative flow cytometric analysis of splenic hCD2^+^TCRβ^+^ T cells from dLckCre*Rosa26^MYC/+^*, dLckCre*Rosa26^MYC/+^Tsc1^fl/fl^*, dLckCre*Rosa26^MYC/+^Slc3a2^fl/fl^*, and dLckCre*Rosa26^MYC/+^Tsc1^fl/fl^Slc3a2^fl/fl^* mice at 13 weeks of age, as well as dLckCre*Rosa26^MYC/+^Tsc1^fl/fl^Slc3a2^fl/fl^* mice at 26 weeks of age. Right panel: Frequencies of splenic hCD2^+^TCRβ^+^ T cells among CD45^+^ cells from dLckCre*Rosa26^MYC/+^* and dLckCre*Rosa26^MYC/+^Tsc1^fl/fl^*mice at 13 weeks of age and hCD2^+^TCRβ^+^CD98^-^ T cells from dLckCre*Rosa26^MYC/+^Slc3a2^fl/fl^*at 13 weeks of age and dLckCre*Rosa26^MYC/+^Tsc1^fl/fl^Slc3a2^fl/fl^*mice at both 13 weeks and 26 weeks of age. (E-H) Representative flow cytometric analysis of MYC (E), phosphorylated S6 (F), and S6 (G) protein as well as FSC-A (H) staining (left panels) and MFI quantification (right panels) in splenic CD4^+^ T cells from wild-type mice, hCD2^+^CD4^+^ T cells from dLckCre*Rosa26^MYC/+^*and dLckCre*Rosa26^MYC/+^Tsc1^fl/fl^* mice, and hCD2^+^CD98^-^CD4^+^ T cells from dLckCre*Rosa26^MYC/+^Slc3a2^fl/fl^* and dLckCre*Rosa26^MYC/+^Tsc1^fl/fl^Slc3a2^fl/fl^* mice at 13 weeks of age. All bar graphs are shown as mean ± SEM, each dot representing one mouse. Data are pooled from multiple experiments. [B, D: one-way ANOVA with Tukey’s multiple comparisons test; C: Log-rank (Mantel Cox) test; E, F, G, H: unpaired t-test, two-tailed, * = p<0.05, ** = p<0.01, *** = p<0.001, **** = p<0.0001, “ns” = not significant.] See also Figure S3.

To explore CD98 function in PTCL lymphomagenesis, we crossed dLckCre*Rosa26^MYC/+^Tsc1^fl/fl^* mice onto the *Slc3a2^fl/fl^*background and confirmed CD98 deletion in dLckCre*Rosa26^MYC/+^Tsc1^fl/fl^Slc3a2^fl/fl^* T cells (Figure S3A). SLC3A2 deficiency prolonged survival of dLckCre*Rosa26^MYC/+^Tsc1^fl/fl^* mice (Figure 3C) in association with time-dependent depletion of PTCL T cells marked by hCD2 expression (Figure 3D). Continuous production of T cells in dLckCre*Rosa26^MYC/+^Tsc1^fl/fl^Slc3a2^fl/fl^* mice might risk the generation of additional mutations, particularly in a lymphopenic environment depleted of SLC3A2-deficient PTCL T cells, to generate escape clones impervious to CD98 deletion (Figure 3C). To bypass this complication, hCD2^+^CD98^-^CD25^-^ T cells from dLckCre*Rosa26^MYC/+^*, dLckCre*Rosa26^MYC/+^Tsc1^fl/fl^*, or dLckCre*Rosa26^MYC/+^Tsc1^fl/fl^Slc3a2^fl/fl^* mice were isolated and transferred to congenically marked wild-type mice that had been sub-lethally irradiated one day prior. While T cells from dLckCre*Rosa26^MYC/+^Tsc1^fl/fl^*mice progressed to become PTCLs causing death of all recipient mice (Figure S3B, S3C), T cells from dLckCre*Rosa26^MYC/+^Tsc1^fl/fl^Slc3a2^fl/fl^*mice were depleted with no pathological consequences in recipient mice (Figure S3B, S3C). Notably, SLC3A2 was dispensable for maintaining T cells that only transcribed *MYC* in dLckCre*Rosa26^MYC/+^Slc3a2^fl/fl^*mice (Figure 3D), implying that PTCL T cells are selectively addicted to CD98 for survival. Further characterization of hCD2^+^ T cells in dLckCre*Rosa26^MYC/+^Tsc1^fl/fl^Slc3a2^fl/fl^*mice prior to their depletion showed that MYC protein expression, mTORC1 signaling, and cell growth all failed to be sustained in the absence of SLC3A2 (Figure 3E-3H). Together, these findings suggest that CD98 is an integral component of the MYC and mTORC1 onco-circuit that enables onco-genotype, i.e. constitutive *MYC* transcription and *Tsc1* deletion, to be manifested as onco-phenotype, i.e. MYC protein overexpression and mTORC1 hyperactivation.

### SLC3A2 elevates mitochondrial membrane potential and curbs ROS accumulation in PTCL

To investigate how PTCL T cells might be addicted to CD98 for survival, we performed additional RNA-sequencing experiments on CD98^-^hCD2^+^CD4^+^CD25^-^ T cells isolated from dLckCre*Rosa26^MYC/+^Tsc1^fl/fl^Slc3a2^fl/fl^*mice and compared their transcriptomes to those of T cells from wild-type and dLckCre*Rosa26^MYC/+^Tsc1^fl/fl^* mice (Table S2). Of note, pathway analyses using HALLMARK and REACTOME gene signatures demonstrated that several mitochondrial metabolic programs including respiratory electron transport, mitochondrial potential-mediated ATP synthesis and heat production, and mitochondrial translation were downregulated in T cells from dLckCre*Rosa26^MYC/+^Tsc1^fl/fl^Slc3a2^fl/fl^* mice relative not only to those from dLckCre*Rosa26^MYC/+^Tsc1^fl/fl^* mice, but also more surprisingly wild-type T cells. (Figure 4A-4C, S4; Table S2).

**Figure 4.**
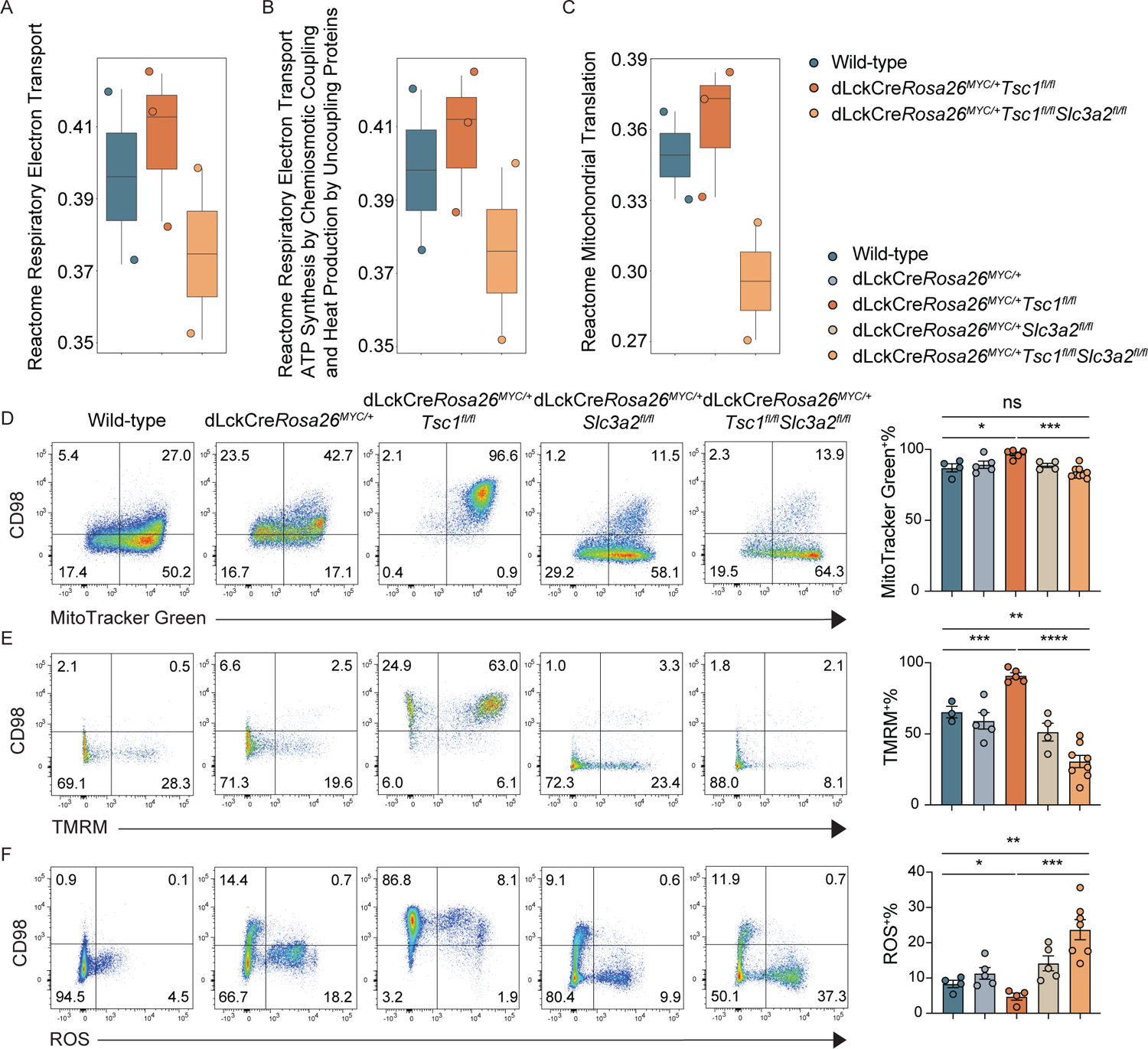
SLC3A2-deficient PTCL cells have defective mitochondrial membrane potential and increased ROS production. (A-C) Box plot showing transcriptional expression (ssGSEA score) of Reactome Respiratory Electron Transport signature (A), Reactome Respiratory Electron Transport ATP Synthesis by Chemiosmotic Coupling and Heat Production by Uncoupling Proteins Pathway Signature (B), and Reactome Mitochondrial Translation Signature (C) in CD4^+^ T cells from wild-type mice, hCD2^+^CD4^+^ T cells from dLckCre*Rosa26^MYC/+^Tsc1^fl/fl^*mice, and hCD2^+^hCD98^-^CD4^+^ T cells from dLckCre*Rosa26^MYC/+^Tsc1^fl/fl^Slc3a2^fl/fl^* mice at 8 weeks of age. (D-F) Representative flow cytometric analysis of CD98 versus MitoTracker Green (D), TMRM (E), and MitoSox Red (F) staining (left panels) and MFI quantification (right panels) in splenic CD4^+^ T cells from wild-type mice and hCD2^+^CD4^+^ T cells from dLckCre*Rosa26^MYC/+^*, dLckCre*Rosa26^MYC/+^Tsc1^fl/fl^*, dLckCre*Rosa26^MYC/+^Slc3a2^fl/fl^*, and dLckCre*Rosa26^MYC/+^Tsc1^fl/fl^Slc3a2^fl/fl^*mice. All bar graphs are shown as mean ± SEM, each dot representing one mouse. Data are pooled from multiple experiments. (D-F: one-way ANOVA with Tukey’s multiple comparisons test, * = p<0.05, ** = p<0.01, *** = p<0.001, **** = p<0.0001, “ns” = not significant.) See also Figure S4.

To determine whether this change in gene expression was functionally consequential, we first measured mitochondrial mass with MitoTracker and found that it was increased in PTCL T cells relative to that of wild-type T cells, but this increase was nullified in the absence of SLC3A2 (Figure 4D). Measurement of mitochondrial membrane potential with TMRM staining showed it to be highest in PTCL T cells, while SLC3A2 deficiency on the PTCL background caused a dramatic decrease to a level even lower than that in wild-type T cells (Figure 4E). Impaired mitochondrial membrane potential led to increased reactive oxygen species (ROS) production in SLC3A2-deficient PTCL T cells, whereas the ROS level was lower in SLC3A2-sufficient PTCL T cells than that in wild-type T cells (Figure 4F). Together, these findings suggest a critical function in enhancing mitochondrial fitness in PTCL T cells for CD98, the lack of which causes decreased mitochondrial membrane potential and increased mitochondrial ROS production that may lead to PTCL demise.

### A low leucine diet disrupts the MYC and mTORC1 onco-circuit in the *MYC/Tsc1* model

Besides assisting assembly of LAT1 and other amino acid transporters, SLC3A2 interacts with β1 integrin to enhance adhesive signaling,^41,42^ and both activities can promote cell survival and proliferation.^43^ Which of these disparate function(s) of SLC3A2 enabled PTCL progression remained unanswered by genetic ablation. Among the imported cargos of LAT1 is the essential amino acid leucine that is not only required for protein synthesis but can also potently stimulate mTORC1 signaling.^44^ To investigate whether the leucine transporter function of SLC3A2 might contribute to PTCL development, we fed dLckCre*Rosa26^MYC/+^Tsc1^fl/fl^* and control mice on an isocaloric low leucine diet (13% of normal chow) starting from 8 weeks of age. The low leucine diet had minimal discernible effect on the health of control mice but delayed the onset of PTCL and greatly enhanced overall survival of dLckCre*Rosa26^MYC/+^Tsc1^fl/fl^* mice (Figure 5A), in association with reduced serum leucine levels (Figure 5B). Supporting this PTCL suppression phenotype, expansion of hCD2^+^TCRβ^+^ lymphoma T cells was inhibited in dLckCre*Rosa26^MYC/+^Tsc1^fl/fl^* mice fed on the low leucine diet (Figure 5C).

**Figure 5.**
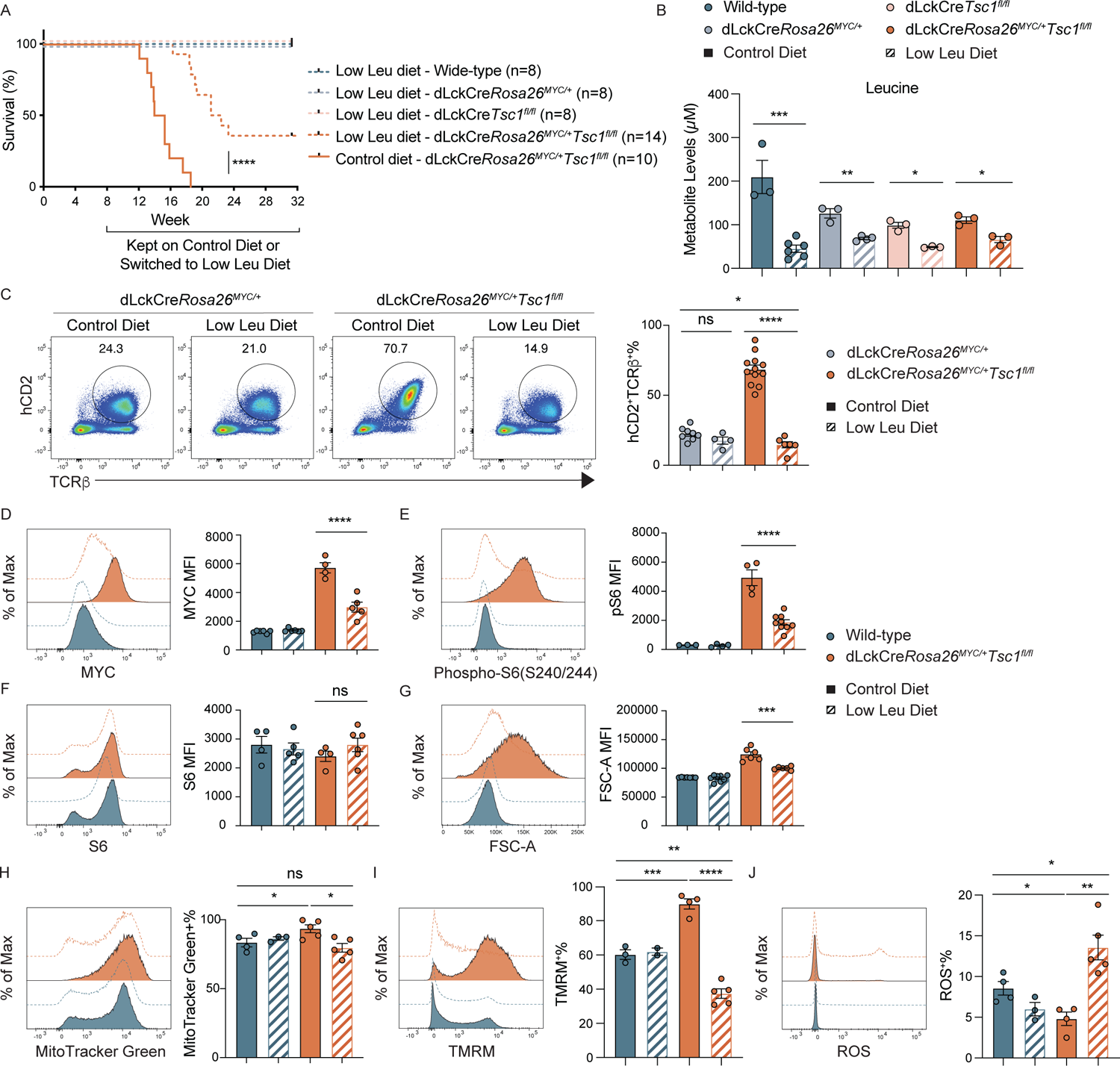
Leucine deprivation disrupts the synergy between constitutive *MYC* transcription and *Tsc1* deletion to interrupt PTCL progression. (A) Survival curves for wild-type, dLckCre*Rosa26^MYC/+^*, dLckCre*Tsc1^fl/fl^*, and dLckCre*Rosa26^MYC/+^Tsc1^fl/fl^* mice switched to an isocaloric low leucine diet (13% leucine of normal chow) starting from 8 weeks of age and dLckCre*Rosa26^MYC/+^Tsc1^fl/fl^* mice kept on normal chow control diet. (B) Serum leucine levels measured by mass spectrometry in wild-type, dLckCre*Rosa26^MYC/+^*, dLckCre*Tsc1^fl/fl^*and dLckCre*Rosa26^MYC/+^Tsc1^fl/fl^* mice fed with control or low leucine diet for 4 weeks. (C) Representative flow cytometric analysis (left panel) and quantification (right panel) of splenic hCD2^+^TCRβ^+^ T cells from dLckCre*Rosa26^MYC/+^*and dLckCre*Rosa26^MYC/+^Tsc1^fl/fl^* mice fed with control or low leucine diet for at least 4 weeks. (D-J) Representative flow cytometric analysis of MYC (D), phosphorylated S6 (E), and S6 (F) protein as well as FSC-A (G), MitoTracker Green (H), TMRM (I), and MitoSox Red (J) staining (left panels) and MFI quantification (right panels) in splenic CD4^+^ T cells from wild-type mice and hCD2^+^CD4^+^ T cells from dLckCre*Rosa26^MYC/+^Tsc1^fl/fl^*mice fed with control or low leucine diet for at least 4 weeks. All bar graphs are shown as mean ± SEM, each dot representing one mouse. Data are pooled from multiple experiments. [A: Log-rank (Mantel Cox) test; B, D, E, F, G: unpaired t-test, two-tailed; C, H, I, J: one-way ANOVA with Tukey’s multiple comparisons test, * = p<0.05, ** = p<0.01, *** = p<0.001, **** = p<0.0001, “ns” = not significant.] See also Figure S5.

Restriction of dietary leucine further reduced MYC protein expression, mTORC1 signaling, and PTCL cell growth in dLckCre*Rosa26^MYC/+^Tsc1^fl/fl^* mice (Figure 5D-5G). Similar to PTCL T cells devoid of SLC3A2, the low leucine diet corrected the increase in mitochondrial mass (Figure 5H) and led to a remarkable loss of mitochondrial membrane potential in PTCL T cells (Figure 5I). Conversely, increased ROS production was detected in PTCL T cells under the low leucine diet condition (Figure 5J). Together, these findings suggest that defective leucine uptake accounts at least in part for the anti-lymphomagenesis phenotypes triggered by SLC3A2 depletion in T cells, and dietary leucine is an integral component of the MYC and mTORC1 onco-circuit operative in dLckCre*Rosa26^MYC/+^Tsc1^fl/fl^* mice.

### Effector T cell responses to infection are highly tolerated to even a leucine-deficient diet

Both MYC and mTORC1 are crucial regulators of metabolic reprogramming and clonal expansion of effector T cells.^39,45^ The profound effects of a low leucine diet on PTCL T cells, but not healthy T cells in the steady state (Figure 5C-5J), raised the question of whether immune challenge-induced effector T cell responses might be similarly sensitive to dietary leucine. To this end, we assessed effector T cell responses in wild-type mice fed on a more stringent isocaloric leucine-deficient (0% of normal chow) diet or normal chow control diet followed by infection with *Listeria monocytogenes* expressing the chicken ovalbumin (LM-OVA) as a model antigen (Figure S5A). Flow cytometry analysis of H-2K^b^-OVA^+^ effector T cells showed that neither the frequency nor the absolute number of bacterial antigen-specific T cells was affected by the leucine-deficient diet (Figure S5B, S5C). Taken together, these findings demonstrate that the detrimental effects of dietary leucine restriction are specific to PTCLs developed in dLckCre*Rosa26^MYC/+^Tsc1^fl/fl^*mice, and do not compromise overall health or normal effector T cell responses to infection.

### Low dietary leucine does not affect tumor development in a *MYC/Depdc5* PTCL model

The profound effect of the low leucine diet on the *MYC/Tsc1* model of PTCL raised a further question as to what extent the impact of reduced leucine uptake on PTCL T cells was caused by defective nutrient mTORC1 signaling versus compromised metabolic function, as leucine is an essential amino acid and cancer cells are metabolically hyperactive. Of note, the nutrient arm of mTORC1 signaling is orchestrated by lysosomal Rag GTPases with cytosolic leucine sensed by the Sestrin family of stress molecules.^44^ Dissociation of leucine from Sestrin1/2 results in their association with the GATOR2 complex and recruitment to the lysosome to inhibit RagA/B as guanine nucleotide inhibitors (GDIs) and permit activation of the GATOR1 complex as a RagA/B GAP.^46^ Of note, GATOR1 together with the KICSTOR complex has a scaffolding function to promote GATOR2 lysosomal localization.^47,48^ Therefore, disruption of the GATOR1 complex can result in constitutive nutrient mTORC1 signaling through multiple mechanisms under nutrient/leucine starvation conditions.

To investigate whether persistent nutrient mTORC1 signaling might, like persistent growth factor signaling, synergize with constitutive *MYC* transcription to induce T cell transformation, we conditionally ablated the GATOR1 component Depdc5 in MYC-expressing T cells by crossing *Depdc5^fl/fl^* mice to the dLckCre*Rosa26^MYC/+^* background. While T cell-specific ablation of Depdc5 did not affect mouse survival, dLckCre*Rosa26^MYC/+^Depdc5^fl/fl^* mice succumbed to death within 20 weeks of age (Figure 6A). Histopathology and flow cytometry analyses revealed extensive T cell expansion and/or infiltration in lymphoid and non-lymphoid organs including spleen, lymph nodes, and liver as well as in blood in dLckCre*Rosa26^MYC/+^Depdc5^fl/fl^*, but not control, mice (Figure S6A, S6B). The expanded T cells had a mixed CD4 and CD8 expression profile in individual mice (Figure S6C) and were monoclonal or oligoclonal (Figure S6D). These findings reveal that constitutive *MYC* transcription and *Depdc5* deletion cooperate to induce PTCL development.

**Figure 6.**
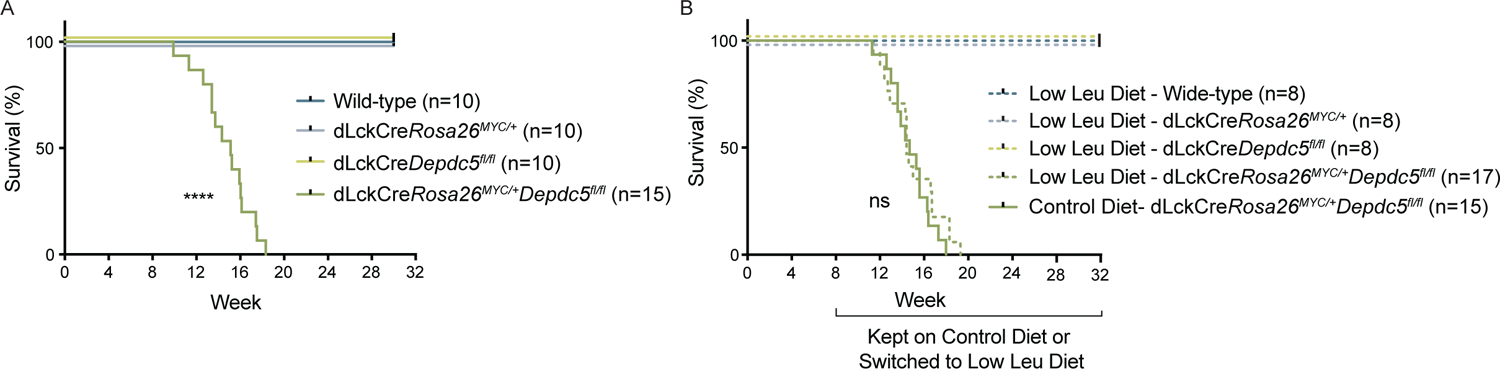
Lethal PTCL driven by constitutive *MYC* transcription and *Depdc5* deletion is unaltered by leucine deprivation. (A) Survival curves for wild-type, dLckCre*Rosa26^MYC/+^*, dLckCre*Depdc5^fl/fl^*, and dLckCre*Rosa26^MYC/+^Depdc5^fl/fl^* mice. (B) Survival curves for wild-type, dLckCre*Rosa26^MYC/+^*, dLckCre*Depdc5^fl/fl^*, and dLckCre*Rosa26^MYC/+^Depdc5^fl/fl^* mice switched to an isocaloric low leucine diet (13% leucine of normal chow) starting from 8 weeks of age and dLckCre*Rosa26*^Myc/+^*Depdc5*^fl/fl^ mice kept on normal chow control diet. [A, B: Log-rank (Mantel Cox) test, “ns” = not significant.] See also Figure S6.

To determine whether PTCLs developed in dLckCre*Rosa26^MYC/+^Depdc5^fl/fl^*mice were sensitive to dietary leucine, we fed dLckCre*Rosa26^MYC/+^Depdc5^fl/fl^* and control mice on an isocaloric low leucine diet (13% of normal chow) starting from 8 weeks of age, which led to reduced serum leucine levels (Figure S6E). However, distinct from the *MYC/Tsc1* model, the low leucine diet did not confer a survival advantage in dLckCre*Rosa26^MYC/+^Depdc5^fl/fl^* mice (Figure 6B) or affect PTCL cell expansion (Figure S6F). In line with these observations, MYC protein overexpression, augmented mTORC1 signaling, and increased PTCL cell size were all comparable between low leucine diet and normal chow control diet groups (Figure S6G). Furthermore, increased mitochondrial mass and hyper-polarized mitochondria membrane potential were unaffected (Figure S6H), while ROS was moderately increased under the low leucine diet condition (Figure S6H). Altogether, these findings suggest that defective nutrient mTORC1 signaling, rather than compromised metabolic function, is the primary reason for low leucine diet suppression of PTCL development in the *MYC/Tsc1* model.

### Low leucine diet demonstrates therapeutic potential for PTCLs in the *MYC/Tsc1* model

Having discovered that the low leucine diet could suppress PTCL development in dLckCre*Rosa26^MYC/+^Tsc1^fl/fl^* mice administered at 8 weeks of age, we wondered whether the low leucine diet might also be effective when used therapeutically in advanced PTCLs. To this end, we isolated PTCL T cells from end-stage dLckCre*Rosa26^MYC/+^Tsc1^fl/fl^*mice and transferred them into congenically marked wild-type recipient mice that had been sub-lethally irradiated one day prior. Recipient mice either remained on the control diet or were transitioned on the same day as PTCL T cell transfer to the low leucine diet. Despite the advanced disease state of the PTCL at the time of transfer we found that the low leucine diet substantially prolonged survival when compared to paired recipients that were kept on control diet (Figure 7A), while the low leucine diet had no effect on overall survival of recipients transferred with wild-type T cells or T cells from dLckCre*Rosa26^MYC/+^* or dLckCre*Tsc1^fl/fl^* mice (Figure 7A). Furthermore, recipient mice transferred with end-stage dLckCre*Rosa26^MYC/+^Depdc5^fl/fl^*donor PTCL T cells were equally susceptible to death under normal chow control diet or the low leucine diet condition (Figure 7B). These findings demonstrate that PTCL T cells developed in dLckCre*Rosa26^MYC/+^Tsc1^fl/fl^* mice are addicted to high concentrations of dietary leucine, which functions as an onco-nutrient to promote lymphoma maintenance. It does so by enabling a MYC/mTORC1 onco-circuit through the nutrient mTORC1 signaling pathway (Figure 7C), a framework of cooperative tumorigenesis with a feedback control module distinct from the conventional oncogene-centric model (Figure S7).

**Figure 7.**
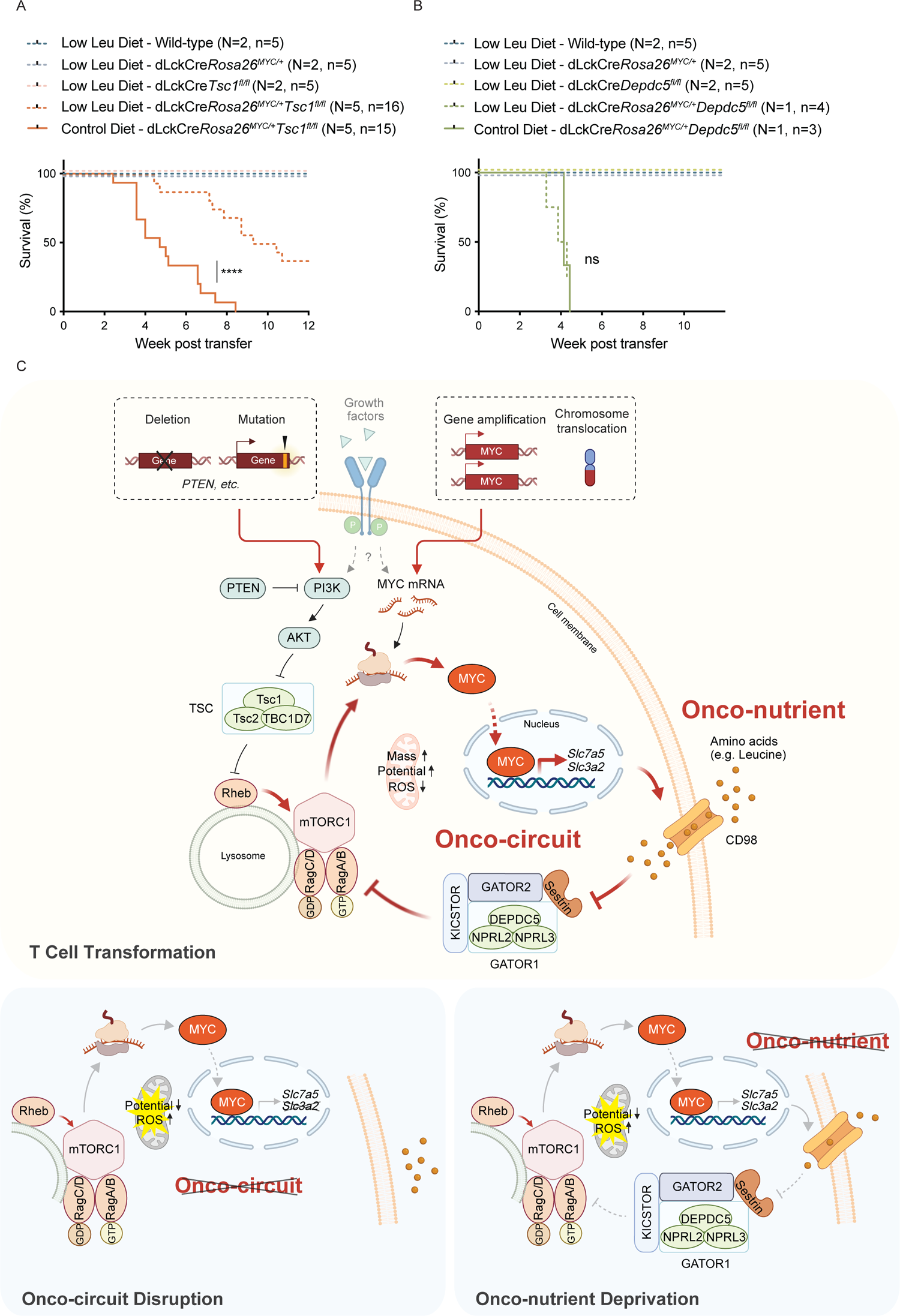
Leucine deprivation improves survival in advanced PTCL in the *MYC/Tsc1* model. (A) Survival curves for sub-lethally irradiated recipient (CD45.1/CD45.2) mice receiving donor (CD45.2) CD4^+^CD25^-^ T cells from wild-type mice (N = 2 donors, n = 5 recipients), hCD2^+^CD4^+^CD25^-^ T cells from dLckCre*Rosa26^MYC/+^* (N = 2 donors, n = 5 recipients) and dLckCre*Tsc1^fl/fl^* (N = 2 donors, n = 5 recipients) mice, and hCD2^+^CD4^+^CD25^-^ PTCL cells from dLckCre*Rosa26^MYC/+^Tsc1^fl/fl^* mice (N = 5 donors, n = 16 recipients) fed on an isocaloric low leucine diet (13% leucine of normal chow) as well as recipient mice fed on normal chow control diet receiving PTCL cells from paired donor dLckCre*Rosa26^MYC/+^Tsc1^fl/fl^*mice (N = 5 donors, n = 15 recipients). Recipient mice were maintained on control diet or switched to low leucine diet starting from the day of cell transfer. (B) Survival curves for sub-lethally irradiated recipient (CD45.1/CD45.2) mice receiving donor (CD45.2) CD4^+^CD25^-^ T cells from wild-type mice (N = 2 donors, n = 5 recipients), hCD2^+^CD4^+^CD25^-^ T cells from dLckCre*Rosa26^MYC/+^* (N = 2 donors, n = 5 recipients) and dLckCre*Depdc5^fl/fl^* (N = 2 donors, n = 5 recipients) mice, and hCD2^+^CD4^+^CD25^-^ PTCL cells from dLckCre*Rosa26^MYC/+^Depdc5^fl/fl^* mice (N = 1 donors, n = 4 recipients) as well as recipient mice fed on normal chow control diet receiving PTCL cells from paired donor dLckCre*Rosa26^MYC/+^Depdc5^fl/fl^*mice (N = 1 donors, n = 3 recipients). Recipient mice were maintained on control diet or switched to low leucine diet starting from the day of cell transfer. (C) Top: Schematic illustrating MYC cooperation with the growth factor arm of mTORC1 signaling to drive T cell transformation. Oncogenic lesions such as *PTEN* deletion frequently target the PI3K/AKT signaling axis to inactivate the Rheb GTPase GAP TSC complex, while the *MYC* oncogene is frequently over-transcribed due to gene amplification or translocation. Among MYC transcriptional targets are *Slc7a5* and *Slc3a2* encoding the heterodimeric large neutral amino acid transporter CD98. CD98-mediated uptake of leucine as an onco-nutrient is sensed via the KICSTOR/GATOR1/GATOR2/Sestrin supercomplex to activate the Rag GTPase complex that recruits mTORC1 to the lysosome surface to be activated by Rheb, which in turn promotes MYC translation in a positive feedback loop. Such aberrant mTORC1 signaling and MYC overexpression synergistically form an onco-circuit to drive PTCL development featuring increased mitochondrial mass, hyperpolarized mitochondrial membrane potential, and reduced mitochondrial ROS accumulation. Bottom: schematics illustrating how the cooperative oncogenic pathway can be disrupted either by genetic ablation of the onco-circuit component *Slc3a2* (left) or deprivation of the key onco-nutrient leucine (right) in association with decreased mitochondrial membrane potential and increased mitochondrial ROS. [A, B: Log-rank (Mantel Cox) test, **** = p<0.0001, “ns” = not significant.] See also Figure S7.

## DISCUSSION

Cancer cells acquire genetic lesions that maladapt signaling, gene expression, and other cellular processes, but how they cooperate to induce and maintain cell transformation is incompletely understood. Using a model of aggressive T cell malignancy triggered by constitutive transcription of *MYC* and deletion of *Tsc1*, we identified a MYC/mTORC1 onco-circuit enabled by an onco-nutrient signaling pathway. While neither a *MYC* nor *Tsc1* genetic lesion (onco-genotype) was sufficient to trigger an effectual functional program (onco-phenotype) manifested in MYC protein expression and mTORC1 signaling, respectively, these functional programs were robustly induced in T cells harboring both. Such MYC/mTORC1 mutual reinforcement was mediated in part by the MYC-induced large neutral amino acid transporter chaperone SLC3A2 and dietary leucine, which, in synergy with *Tsc1* deletion, triggered robust mTORC1 activation. In turn, mTORC1 promoted mitochondrial fitness and MYC translation, forming a positive feedback onco-circuit. Of note, elevated leucine mTORC1 signaling fostered not only induction but also maintenance of *MYC/Tsc1* T cell malignancy, yet leucine restriction did not impair microbial antigen-specific T cell responses. These findings reveal that *MYC/Tsc1* lymphoma T cells, but not healthy T cells, are addicted to persistent nutrient mTORC1 signaling, and this aspect of onco-circuit addiction features hybrid characteristics of non-oncogene addiction^4^ and oncogene addiction^3^. Specifically, similar to non-oncogene addiction, the pathway upon which the onco-circuit relies—LAT1-mediated leucine acquisition—need not harbor mutations, but like oncogene addiction, the pathway is directly oncogenic as leucine promotes mTORC1 signaling and MYC protein expression.

The finding that T cell lymphoma manifestation requires interplay between constitutive *MYC* transcription and *Tsc1* loss with support of transporter-mediated uptake of dietary leucine suggests it is the dynamic positive feed-back circuit, and not merely static oncogenic lesions, to which the cancer cells are addicted. Given the pleiotropic effects and frequent dysregulation of MYC and mTORC1 in T cell lymphomas, additional genes and pathways may further contribute to this MYC/mTORC1 onco-circuit. Notably, a recent study revealed malic enzyme 2 (ME2) to be a direct transcriptional target of MYC essential for MYC-induced T cell lymphomagenesis.^49^ ME2 is required to maintain redox homeostasis and stimulate mTORC1 by modulating glutamine metabolism, with the latter potentially because of heightened LAT1-mediated glutamine efflux in exchange for increased leucine uptake.^50^ Speculatively, the longer latency period in that model prior to lymphomagenesis may be due to selection for mutations that functionally inactivate TSC to instigate a MYC/mTORC1 onco-circuit similar to that in the *MYC/Tsc1* model, with the MYC-ME2 module acting in concert with the MYC-LAT1 module in support of continuous leucine uptake to sustain malignancy.

Various forms of cancer cell addiction to amino acids, and ensuing pharmacologic and dietary therapeutic strategies, have been described in tumor models or clinical practice.^51^ While diverging in important ways, these strategies all posit primarily metabolic interpretations for differential cancer cell sensitivity to amino acid restriction. For example, the original and to-date most successful clinical application of an amino acid-depleting therapy is the use of L-asparaginase in T-cell acute lymphoblastic leukemia (ALL), as ALL cells are often auxotrophic for the otherwise non-essential amino acid asparagine.^52^ Amino acids involved in one-carbon metabolism including methionine,^53^ glycine,^54^ and serine^55^ are required for synthesis of not only proteins but also nucleotides, phosphatidylserine, and other biomolecules without which cancer cells cannot proliferate. Similarly, depletion of glutamine^56^ and cysteine^57^ can also be limiting for NADPH and glutathione production and thus redox homeostasis, with deficiencies leading to cell death including by ferroptosis.

In many of these examples, mTORC1 activation has also been noted to accompany elevated amino acid uptake, and mTORC1 inhibition can synergize with amino acid restriction.^38,58,59^ However, disentangling the relative contributions of metabolic versus signaling functions for amino acid acquisition has remained elusive. To approach this long-standing question, we compared leucine sensitivity in the *MYC/Tsc1* model with the *MYC/Depdc5* model, in which leucine is largely dispensable for mTORC1 activation but remains essential for biosynthesis. Surprisingly, a low though non-null level of dietary leucine had a profound impact on mouse survival and cancer cell phenotype in the *MYC/Tsc1* model but not *MYC/Depdc5* model. This finding strongly supports signaling primacy for leucine in the MYC/mTORC1 onco-circuit, a role we refer to as that of an onco-nutrient. It is unlikely that amino acid sensitivity for purposes of signaling greatly surpasses that of metabolism in most cellular contexts. For instance, the binding affinity (Kd) of leucine to the mTORC1 signaling sensor Sestrin 2 is around 20 μM,^60^ while the leucyl-tRNA synthetase Michaelis constant (*KM*) for leucine activation is around 45 μM^61^. Increased sensitivity of lymphoma T cells to leucine mTORC1 signaling thus may be due to increased dependence of metabolically supercharged cancer cells on persistent mTORC1 activation to proactively serve as a buffer for particularly variable nutrient availability. This may be accomplished through the MYC/mTORC1 onco-circuit itself, which induces high levels of LAT1 expression to accommodate low levels of circulating leucine. This crucial component of the circuit is disrupted in the *MYC/Tsc1* model but not *MYC/Depdc5* model, and though leucine’s primary role in maintaining it occurs through mTORC1 signaling, a secondary result may be further inadequate nutrient acquisition and metabolic insufficiency consequent to failed induction of LAT1. To this end, the lack of effect of a leucine-deprived diet on physiological T cell responses to infection is also noteworthy, considering that deletion or inactivation of LAT1, mTORC1, or mTORC1 and c-MYC cross-regulation during asymmetric division of activated T cells affects proliferation and differentiation of effector T cells.^62–67^ It is possible that the selectively deleterious effect of leucine starvation on lymphoma cells is due to metabolic amplification consequent to uncontrolled oncogenic MYC expression, while endogenous c-MYC expression during physiological T cell responses is dynamically regulated and can effectively adapt to environmental signals such as dietary leucine that regulate mTORC1 activation.

Both genetic and dietary restriction of cytosolic leucine resulted in severe mitochondrial dysfunction in *MYC/Tsc1* lymphoma T cells, marked by decreased mass and membrane potential and increased ROS. This decline in mitochondrial fitness may represent an active sensing mechanism, wherein decoupled MYC and mTORC1 signaling is conveyed to the mitochondria to initiate the cell death process. In support of this possibility, mTORC1 has been shown to promote translation of the mitochondrial anti-apoptotic protein Mcl-1,^68^ while its inhibition triggers upregulation of the proapoptotic protein BMF in MYC-driven B cell lymphoma models.^69^ Furthermore, mTORC1 may regulate multiple aspects of mitochondrial biogenesis and activity, including via translation of mitochondrial ribosomal proteins,^70^ which we noted to be decreased in the setting of SLC3A2 depletion in *MYC/Tsc1* lymphoma T cells, albeit at the transcriptional level. Alternatively, the observed phenotype may be secondary to a cell death cascade initiated on account of overwhelmed stress response mechanisms and thus represent a passive process.^71^ The lack of clarity on the precise causal chain leading from diminished nutrient signaling to diminished mitochondrial fitness will merit further investigation. In either case, elucidation of a MYC/mTORC1 onco-circuit and its dependence on nutrient acquisition—particularly given the wide therapeutic window afforded by differential leucine requirements for *MYC/Tsc1* T cell lymphomagenesis relative to infection response—will motivate therapeutic exploration in the most aggressive, MYC/mTORC1-coupled peripheral T cell lymphoma cases. Along with dietary restriction of leucine, this may be approached using existing clinical-stage LAT1-blocking inhibitors.^72,73^

Beyond lymphomas, *MYC* is dysregulated in most solid tumors to the point that MYC signaling is considered a molecular hallmark of cancer.^74^ Likewise, many oncogenic pathways such as PI3K converge on the TSC complex, rendering growth factor-independent activation of this branch of mTORC1 signaling another prominent cancer hallmark.^75^ Of note, analysis of somatic copy number alterations (SCNAs) in chromosomal instability (CIN)-driven tumors such as high-grade serous ovarian carcinoma, squamous lung cancer, and *TP53*-mutant triple negative breast cancer revealed frequent co-occurrence of amplification or gain of the *MYC* gene and genes of the PI3K pathways such as *PIK3CA*,^76^ suggesting their functional cooperation in cancer progression. Indeed, we have recently shown that constitutive *MYC* transcription synergizes with PI3K-activating polyomavirus middle tumor antigen (PyMT) oncoprotein to induce fulminant tumor growth in a transgenic model for aggressive breast cancer.^77^ MYC-overexpressing PyMT cancer cells acquire a super-competitor cell differentiation state that is dependent on their superlative activation of mTORC1.^77^ While the function of transporter (e.g. LAT1)-mediated nutrient acquisition in mTORC1 activation remains to be determined, engulfment-mediated nutrient acquisition via the endolysosomal lipid kinase PIKfyve is essential for mTORC1 signaling and survival of MYC-overexpressing PyMT cancer cells.^77^ These findings suggest that addiction to a nutrient MYC/mTORC1 onco-circuit may be a general feature of aggressive cases across cancer types, although the exact enabling mechanism may be context-dependent in terms of the route of nutrient acquisition and sensing mechanism. Such an intimate coupling of MYC and mTORC1 activity is likely rooted in its fundamental role in cell fitness control throughout mammalian biology, including as early as embryonic stem cell fate specification.^78^

In summary, in a model of aggressive peripheral T cell lymphoma, we have herein discovered a prominent mechanism by which oncogenic genetic lesions cooperate to induce and sustain malignancy via a MYC/mTORC1 onco-circuit. High and specific sensitivity of cancer cells to leucine as an onco-nutrient motivates development of cancer therapeutics targeting the nutrient mTORC1 signaling pathway, including additional non-mutated components of the onco-circuit. The framework of cooperative tumorigenesis described in this study should also motivate additional scrutiny into how onco-phenotype is manifested from onco-genotype for other complex cancer drivers, and how the circuits engaged in this process engender addiction and thus vulnerability.

## ACKNOWLEDGEMENTS

We thank members of the M.O.L. laboratory for helpful discussions. We acknowledge the use of the Integrated Genomics Operation Core, funded by the NCI Cancer Center Support Grant (CCSG, P30 CA08748), Cycle for Survival, the Marie-Josée and Henry R. Kravis Center for Molecular Oncology, and the Center of Comparative Medicine & Pathology at Memorial Sloan Kettering Cancer Center. This work was supported by NIAID (R01 AI102888) and a Functional Genomics Initiative Grant to M.O.L, and NIH (P01 HL-151433) to M.H.G.

## AUTHOR CONTRIBUTIONS

X.W. and M.O.L. were involved in all aspects of this study, including planning and performing experiments, analysis and interpretation of data, and writing the manuscript. A.E.C. was involved in planning and analyzing sequencing and other experiments and writing the manuscript. M.H.D. made a relevant initial observation that led to this project. J.S.B. and L.W.S.F. performed serum leucine measurements. T.W.H., Z.X., I.M., C.E., K.J.C., and X.Z. assisted with mouse colony management and performed experiments. M.H.G. provided a key mouse line. M.S.L and S.M.H provided guidance related to human PTCL. All authors provided critical scientific feedback on the results and manuscript.

## STAR METHODS

### Lead Contact and Materials Availability

Further information and requests for resources and reagents should be directed to and will be fulfilled by the lead contact, Dr. Ming O. Li. The raw sequencing data generated in this study will be deposited to GEO upon publication.

### Animal Experimental Model Details

CD4Cre, *Rosa26^MYC/+^*, *Igs2^MYC(T58A)/+^*, *Tsc1^fl/fl^*, dLckCre, CD45.1 and CD45.2 mice were purchased from Jackson Laboratory. *Depdc5^fl/fl^*mice were generated in the laboratory of Dr. Ming O. Li and previously described.^77^ *Slc3a2^fl/fl^* mice were previously described.^43^ All mice used in these studies were on the C57BL/6 background. Mice were maintained under specific pathogen-free conditions, and animal experimentation was conducted in accordance with procedures approved by the Institutional Animal Care and Use Committee of Memorial Sloan Kettering Cancer Center. Listeria monocytogenes-OVA infection conformed to all relevant regulatory standards. In dietary switch experiments, irradiated isocaloric low Leucine (5G7M, TestDiet) diet or 0% Leucine (5WH9, TestDiet) diet were substituted for standard chow for mice. Mice aged between 8-13 weeks were used for experiments unless otherwise noted. No sex differences in phenotype were noted.

### Method Details

#### Immune cells isolation

Immune cells were isolated from mouse thymus, spleen, peripheral (axillary, brachial, inguinal and superficial cervical) lymph nodes, liver and blood. For thymus, spleen, and peripheral lymph nodes, single cell suspension was obtained by grinding tissues and filtered through a 70 μm cell strainer. Cells were then depleted of erythrocytes by hypotonic lysis for spleen and blood. For liver, tissues were manually dissociated using a razor blade and digested with 280 U/ml Collagenase Type III and 4 μg/ml DNase I in HBSS for 30 min at 37 °C with periodic vortex. Digested tissues were then filtered through a 70 μm cell strainer and pelleted. To isolate immune cells, pellets were resuspended in 44% Percoll and layered above 66% Percoll. Samples were centrifuged at 1,900 *g* for 30 min without brake. Cells at the interface were collected, stained and analyzed by flow cytometry or sorting.

#### Mouse necropsy phenotyping

Following euthanasia, blood was collected in anticoagulant-coated tubes for Wright Giemsa stain. Tissues were harvested and fixed in 10% neutral-buffered formalin and embedded in paraffin. Tissues were sectioned at 5 μm and stained with hematoxylin & eosin (H&E). H&E slides and blood smears were examined by a board-certified veterinary pathologist.

#### Flow cytometry

Cells were pre-stained with Ghost Dye (Tonbo) at 4 °C for 10 min for the exclusion of dead cells. Cells were then stained with panels of antibodies for 30 min on ice in 1× PBS containing 2% FBS and 2 mM EDTA FACS buffer. For intracellular staining, Tonbo transcription factor staining kit was used to fix and permeabilize cells for 30 min on ice, followed by intracellular staining for 30 min on ice. For cytokine staining, cells were stimulated with 50 ng/ml phorbol 12-myristate 13-acetate, 1 mM ionomycin and GolgiStop for 4 hours prior to staining. For MYC protein, phosphorylated S6 and S6 staining, spleens were isolated and immediately meshed into 4% paraformaldehyde solution for 15 min at room temperature, followed by 1-hour permeabilization with Tonbo fix/perm buffer at 4 °C. Cells were then washed and incubated with primary antibodies for 1 hour at room temperature, then secondary antibodies for 45 min at 4 °C, followed by flow cytometry analysis. All samples were acquired with Fortessa (Becton Dickson) and analyzed with FlowJo software (TreeStar).

#### Cell sorting

After gating on morphology and singlets, splenic alive CD45^+^ cells were further gated on TCRβ^+^CD25^-^ and hCD2^+^ (if from mice with *Rosa26^MYC/+^* allele) and CD98^-^ (if from mice with *Slc3a2*^fl/fl^ allele) as total T cells used in cell transfer experiments into sub-lethally irradiated recipients. TCRβ^+^CD25^-^(hCD2^+^)(CD98^-^) cells were further gated on CD4^+^ T cells for bulk RNA sequencing. Cell sorting was conducted on Aria II (Becton Dickson) using a 70 μm nozzle controlled by software FACSDiva (Becton Dickson).

#### Adoptive cell transfer

Splenic TCRβ^+^CD25^-^(hCD2^+^)(CD98^-^) total T cells (CD45.2) were FACS-sorted. 1×10^6^ sorted cells were transferred intravenously into 8-10 weeks old CD45.1/CD45.2 wild-type mice that had been sub-lethally irradiated with 600 Gy the day before. In dietary switch experiment, recipient mice were switched to low leucine diet on the day of adoptive cell transfer. Recipient mice were monitored for survival continuously.

#### Mitochondrial mass, mitochondrial ROS staining

1×10^6^ splenocytes were stained with live/dead stain and cell surface markers first at 4 °C for 30 min. Cells were then washed with pre-warmed HBSS once and incubated with 5 μM MitoSoxRed at 37 °C for 10 min to measure mitochondrial ROS, or with 100 nM of MitoTracker Green at 37° C for 30 min to measure mitochondrial mass. After incubation, cells were washed twice with FACS buffer and analyzed by flow cytometry immediately.

#### Mitochondrial cell membrane potential measurement

1=10^6^ splenocytes were stained with live/dead stain and cell surface markers first at 4 °C for 30 min. Cells were then washed with pre-warmed PBS once and incubated with 20 nM TMRM at 37 °C for 30 min in polypropylene tubes to measure mitochondrial membrane potential. After incubation, cells were washed once with PBS and analyzed by flow cytometry immediately.

#### Serum leucine concentration measurement

50 μl of blood was collected from tail of each mouse fed for at least 4 weeks on low leucine or control diet, coagulated on ice for 30 min and spun at 2,000 *g* for serum collection at 4°C. 10 μl serum sample was transferred to a new tube to which 990 μl ice-cold 80% methanol supplemented with 2 μM deuterated D-2-hydroxyglutaric-2,3,3,4,4-d5 acid (d5-2HG) was added. Samples were incubated overnight at −20 °C to extract metabolites. The next day, samples were spun down at 4 °C, 21,000 *g* for 20 min and dried in an evaporator (Genevac EZ-2 Elite). Dried pellets were resuspended in 40 mg/ml methoxyamine hydrochloride in pyridine by shaking for 2 hours at 30 °C. Samples were derivatized with 80 μl of N-methyl-N-(trimethylsilyl) trifluoroacetamide and 70 μl ethyl acetate and incubated for 30 min at 37 °C. Analysis of samples was performed using an Agilent 7890B Gas Chromatograph coupled to Agilent 5977B mass selective detector in splitless mode using constant helium gas flow at 1 ml/min. 1 μl of derivatized metabolites was injected onto an HP-5ms column and the gas chromatograph oven temperature ramped up from 60 °C to 290 °C over 25 min. Peaks were assessed with MassHunter software v.B.08 (Agilent) and normalized to internal standards (d5-2HG). Absolute concentrations of serum leucine were calculated by including a standard curve of known leucine concentrations. Leucine levels were derived by quantifying the following ions: d5-2HG, 354 m/z; Leucine, 260 m/z. All of the peaks were manually inspected and verified relative to known spectra for each metabolite.

#### *Listeria Monocytogenes-*OVA (LM-OVA) infection

Frozen aliquot/vial of LM-OVA (stored at −80 °C) was thawed and 200 μl of bacteria was added into 5 ml of sterile brain heart infusion (BHI) media. Bacterial culture was incubated 3.5 to 4 hours at 37 °C shaking at 225 rpm. Bacterial OD600 was monitored and ready when reached between 0.1-0.2 (OD600 = 0.1 is equal to 1×10^8^ bacteria/ml). Wild-type mice were intravenously injected with 5×10^3^ colony-forming units (CFU) of LM-OVA. Antigen specific effector CD8^+^ T cell responses were assessed on day 8.

#### Human Microarray data

For human PTCL microarray data, a total of 372 samples (after excluding cell line controls) as well as clinical annotations were downloaded directly from GEO from a total of four series/publications: GSE14879,^79^ GSE19069,^80^ GSE58445,^20^ and GSE6338.^81^

#### RNA sequencing

For bulk RNA sequencing, splenic T cells were sorted into Trizol LS and flash frozen. Phase separation in cells lysed in TRIzol was induced with 200 μl chloroform and RNA was extracted from the aqueous phase using the MagMAX *mir*Vana Total RNA Isolation Kit on the KingFisher Flex Magnetic Particle Processor (Thermo Scientific) according to the manufacturer’s protocol with 350 μl input. Samples were eluted in 33 μl elution buffer. After RiboGreen quantification and quality control by Agilent BioAnalyzer, 2 ng total RNA with RNA integrity numbers ranging from 7.7 to 10 underwent amplification using the SMART-Seq v4 Ultra Low Input RNA Kit (Clonetech catalog # 63488), with 12 cycles of amplification. Subsequently, 10 ng of amplified cDNA was used to prepare libraries with the KAPA Hyper Prep Kit (Kapa Biosystems KK8504) using 8 cycles of PCR. Samples were barcoded and run on a NovaSeq 6000 in a PE100 run, using the NovaSeq 6000 S4 Reagent Kit (200 cycles) (Illumina). An average of 62 million paired reads were generated per sample, the percent of mRNA bases per sample ranged from 57% to 93%, and ribosomal reads averaged 0.4%.

### Quantification and Statistical Analysis

Paired-end reads were downloaded in FASTQ format from 21 total samples comprising the following combinations of genotypes and disease time points: wild-type (n=2), CD4Cre*Tsc1^fl/fl^* (n=2), CD4Cre*Rosa26^MYC/+^* (n=2), CD4Cre*Rosa26^MYC/+^Tsc1^fl/fl^* (n=3), dLckCre*Rosa26^MYC/+^Tsc1^fl/fl^* (n=3) at an early time point (8 weeks), dLckCre*Rosa26^MYC/+^Tsc1^fl/fl^*(n=3) at a late time point (13 weeks), dLckCre*Rosa26^MYC/+^Depdc5^fl/fl^*(n=3) at a late time point (13 weeks) and dLckCre*Rosa26^MYC/+^Tsc1^fl/fl^Slc3a2^fl/fl^* (n=3) at an early time point (8 weeks). Samples were quantified at the transcript level with Kallisto (v0.46.0) (Bray et al., 2016) using the mm10 reference and default parameters, and imported into a gene X sample count matrix in R using the “tximport” package.^82^

For pairwise differential analysis related to Figures 2B, S2C, 3A, 4A-C, and S4: Pairwise differential analyses between select genotypes/timepoints of interest was conducted using the “DESeq2” package,^83^ and genes were included in the appended lists of differentially expressed genes (Tables S1 and S2) if they met all three of the following criteria: base mean expression > 100, p-value < 0.01, and abs(log2 fold change) > 1. The MYC-mTORC1 Mouse PTCL Signature used for investigation of human samples was composed of differentially expressed genes defined in this manner and up in dLckCre*Rosa26^MYC/+^Tsc1^fl/fl^* (late) versus wild-type mice. Enriched gene signatures between conditions were identified using pre-ranked GSEA (ranked by log fold change from DESeq2 differential expression analysis) as implemented by the R package “clusterProfiler”.^84^ Pre-defined signatures tested for enrichment included all REACTOME and HALLMARK canonical pathways in MSigDB v7.1,^85^ and were included in lists of enriched signatures if Benjamin-Hochberg FDR P-value was below 0.01 and abs(NES) > 1.75 (where NES is normalized enrichment score). Signatures shown in heatmaps were hand-picked from this list of enriched signatures to minimize functional redundancy across canonical pathways and displayed by their Z-Score across samples included in the heatmap.

For principal components analysis (PCA) related to Figure S2B: variance-stabilized transformed values (computed with DESeq2) from the top 500 most variable genes were included as input, and the first two principal components are shown.

For TCR clonality analysis related to Figure 2A, S2A, and S6D: To calculate degree of clonal expansion of lymphoma clones, TCRβ CDR3 region sequences were extracted using the MiXCR software package.^34^ The clonality of the resultant TCR repertoire was then computed as 1 - the normalized Shannon Entropy of the repertoire, with entropy calculated using the observed proportion of reads mapping to each unique TCRβ CDR3 sequence. This also correlated with clonality calculated using TRBV-mapped reads with Pearson correlation coefficient 0.97. Student’s *t*-test was used to assess for the difference in clonality between conditions.

For human microarray analysis: Meta data was extracted from GEO using the getGEO() function from the GEOquery R package.^86^ Raw data was downloaded in the CEL format using the affy R package,^87^ and normalization was done using the robust multi-array (RMA) technique.^88^ Given disparate sources of data batch correction was performed using empirical Bayes methods as implemented in the R package ComBat.^89^ For genes with more than one probe, the probe with the highest median absolute deviation was used for all downstream analyses.

Related to Figure 2G: Expression of the murine PTCL gene signature, as well as MSigDB REACTOME and HALLMARK canonical pathways, was calculated in each human sample using Single-sample Gene Set Enrichment Analysis (ssGSEA) via the R package “GSVA”.^90^ For Kaplan-Meier survival analysis related to Figure 2H, samples were divided into those above and below the median ssGSEA expression of the murine PTCL signature, and survival analysis was done using the R package “survfit”.^100^

All bar graphs are displayed as mean ± SEM, each dot representing one mouse. Data are pooled from multiple experiments. For survival analysis, Log-rank (Mantel Cox) test was performed using GraphPad Prism software. For pair-wise comparisons, for two samples, unpaired student t test, two-tailed was conducted, and for three or more samples, one-way ANOVA with Tukey’s test for corrections was conducted using GraphPad Prism software. * = p<0.05, ** = p<0.01, *** = p<0.001, **** = p<0.0001, “ns” = not significant.

## Data and Code Availability

The datasets supporting the current study are available from the corresponding author on request.

## KEY RESOURCE TABLE

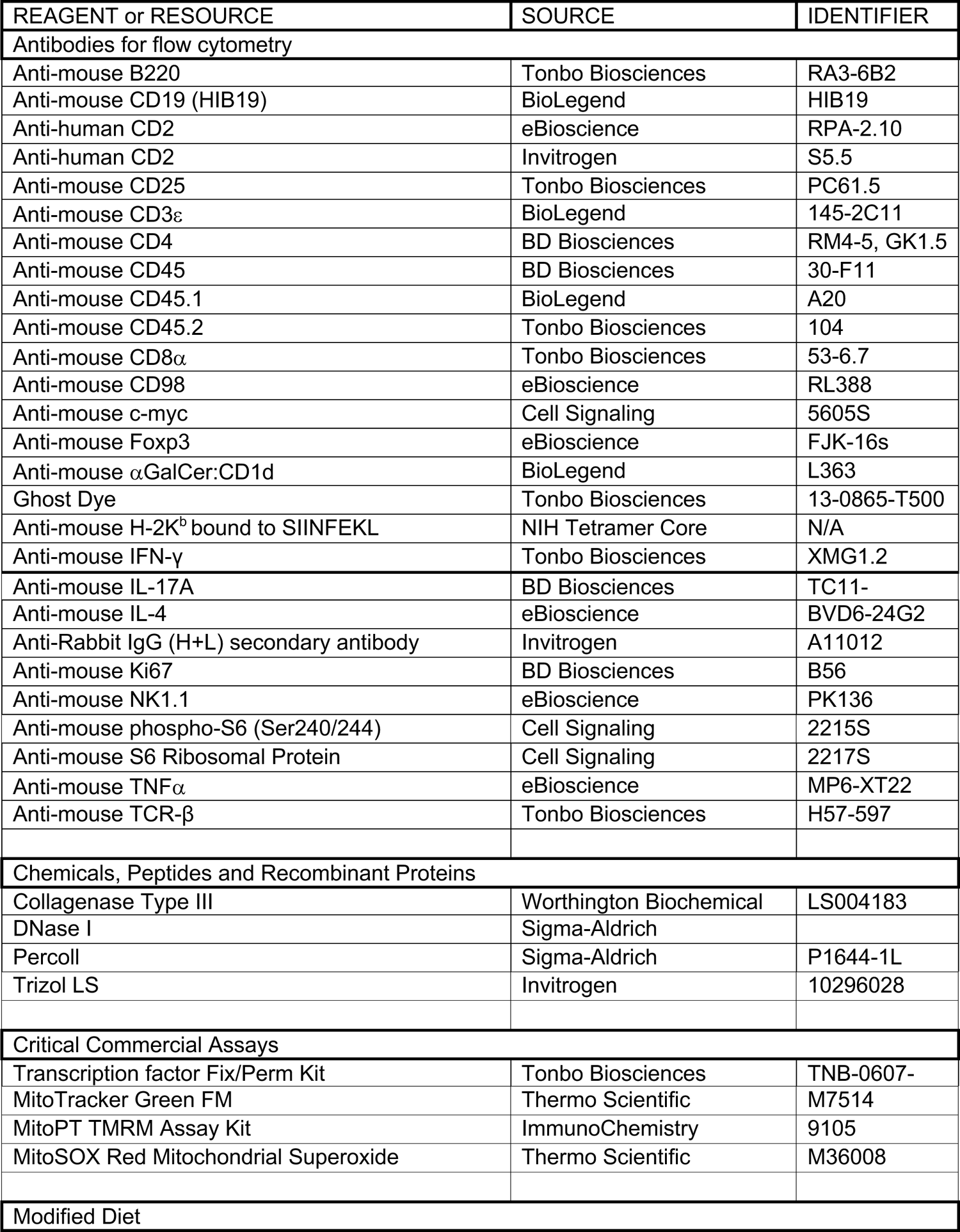
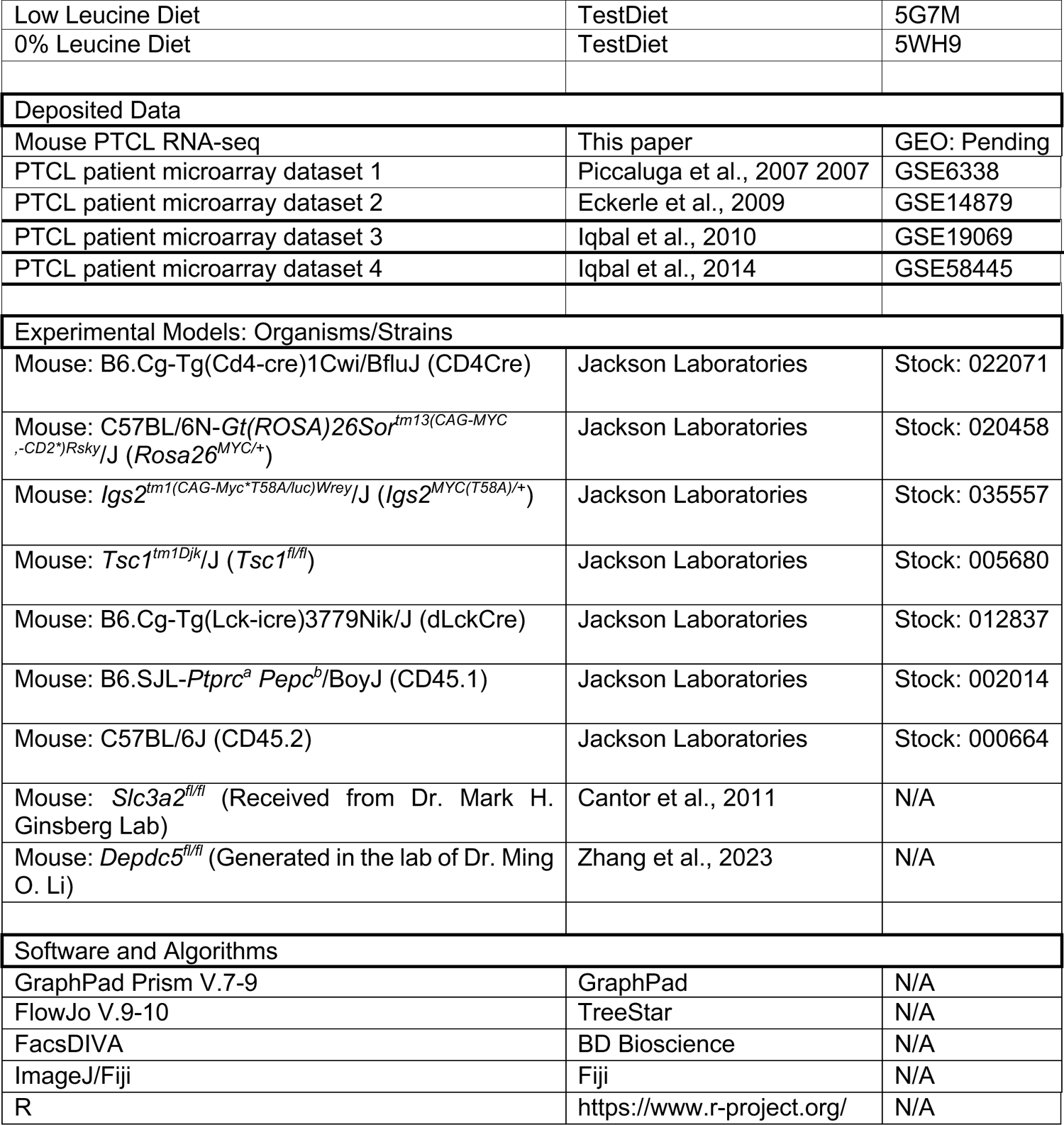

### SUPPLEMENTAL INFORMATION TITLES AND LEGENDS

**Figure S1.**
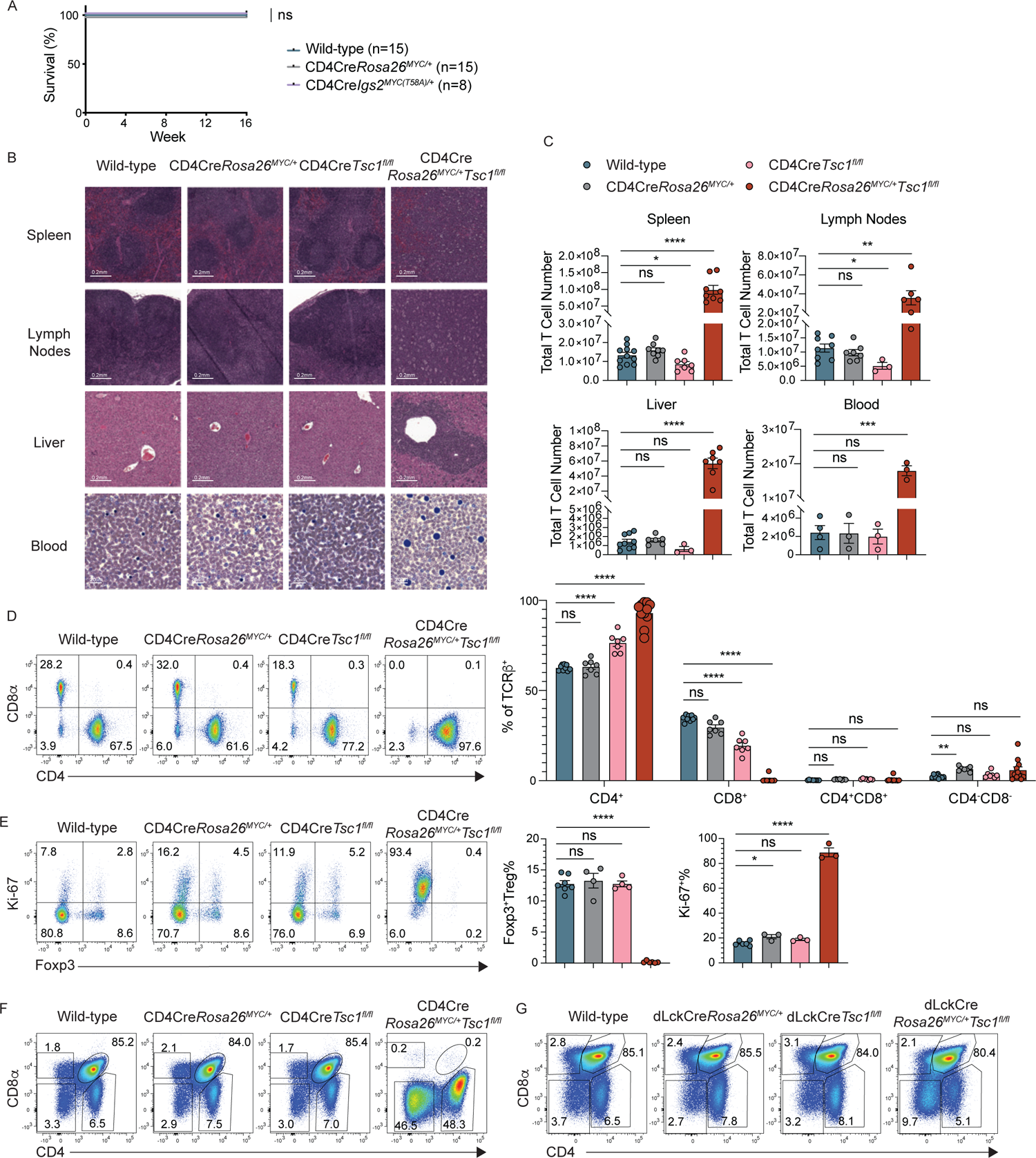
CD4Cre-driven constitutive *MYC* transcription synergizes with *Tsc1* deletion in T cells to trigger a lethal phenotype in mice. Related to Figure 1. (A) Survival curves for wild-type, CD4Cre*Rosa26^MYC/+^*, and CD4Cre*Igs2^MYC(T58A)/+^* mice. (B) Rows 1-3: Representative images of hematoxylin and eosin (H&E) staining of spleen, lymph node and liver in 4-week-old wild-type, CD4Cre*Rosa26^MYC/+^*, CD4Cre*Tsc1^fl/fl^*, and CD4Cre*Rosa26^MYC/+^Tsc1^fl/fl^*mice. Original magnification, 21x. Scale bar 0.2mm. Row 4: Representative images of Giemsa stain of blood from 4-week-old wild-type, CD4Cre*Rosa26^MYC/+^*, CD4Cre*Tsc1^fl/fl^*, and CD4Cre*Rosa26^MYC/+^Tsc1^fl/fl^* mice. Original magnification, 63x. Scale bar 20μm. (C) Quantification of total T cell numbers in spleen, lymph nodes, liver and blood from wild-type, CD4Cre*Rosa26^MYC/+^*, CD4Cre*Tsc1^fl/fl^*, and CD4Cre*Rosa26^MYC/+^Tsc1^fl/fl^*mice. (D) Representative flow cytometric analysis of CD4 and CD8α expression (left panel) in splenic T cells from wild-type, CD4Cre*Rosa26^MYC/+^*, CD4Cre*Tsc1^fl/fl^*, and CD4Cre*Rosa26^MYC/+^Tsc1^fl/fl^*mice at 4 weeks of age. Right panel: Frequencies of CD4^+^, CD8^+^, CD4^+^CD8^+^ and CD4^-^CD8^-^ splenic T cells (right panel) from wild-type, CD4Cre*Rosa26^MYC/+^*, CD4Cre*Tsc1^fl/fl^*, and CD4Cre*Rosa26^MYC/+^Tsc1^fl/fl^*mice. Enlarged dots represent statistics of expanded T cell populations. (E) Representative flow cytometric analysis of Ki-67 and Foxp3 expression (left panel) in splenic CD4^+^ T cells from wild-type, CD4Cre*Rosa26^MYC/+^*, CD4Cre*Tsc1^fl/fl^*, and CD4Cre*Rosa26^MYC/+^Tsc1^fl/fl^*mice at 4 weeks of age. Frequencies of Foxp3^+^ Treg cells (middle panel) and Ki-67^+^ cells (right panel) among splenic CD4^+^ T cells from wild-type, CD4Cre*Rosa26^MYC/+^*, CD4Cre*Tsc1^fl/fl^*, and CD4Cre*Rosa26^MYC/+^Tsc1^fl/fl^* mice at 4 weeks of age. (F) Representative flow cytometric analysis of CD4 and CD8α expression in the thymus from 4-week-old wild-type, CD4Cre*Rosa26^MYC/+^*, CD4Cre*Tsc1^fl/fl^*, and CD4Cre*Rosa26^MYC/+^Tsc1^fl/fl^*mice. Data are representative of 4 independent experiments. (G) Representative flow cytometric analysis of CD4 and CD8α expression in the thymus from 7-week-old wild-type, dLckCre*Rosa26^MYC/+^*, dLckCre*Tsc1^fl/fl^*, and dLckCre*Rosa26^MYC/+^Tsc1^fl/fl^* mice. Data are representative of 7 independent experiments. All bar graphs are shown as mean ± SEM, each dot representing one mouse. Data are pooled from multiple experiments. [A: Log-rank (Mantel Cox) test; C, D, E: one-way ANOVA with Tukey’s multiple comparisons test, * = p<0.05, ** = p<0.01, *** = p<0.001, **** = p<0.0001, “ns” = not significant.]

**Figure S2.**
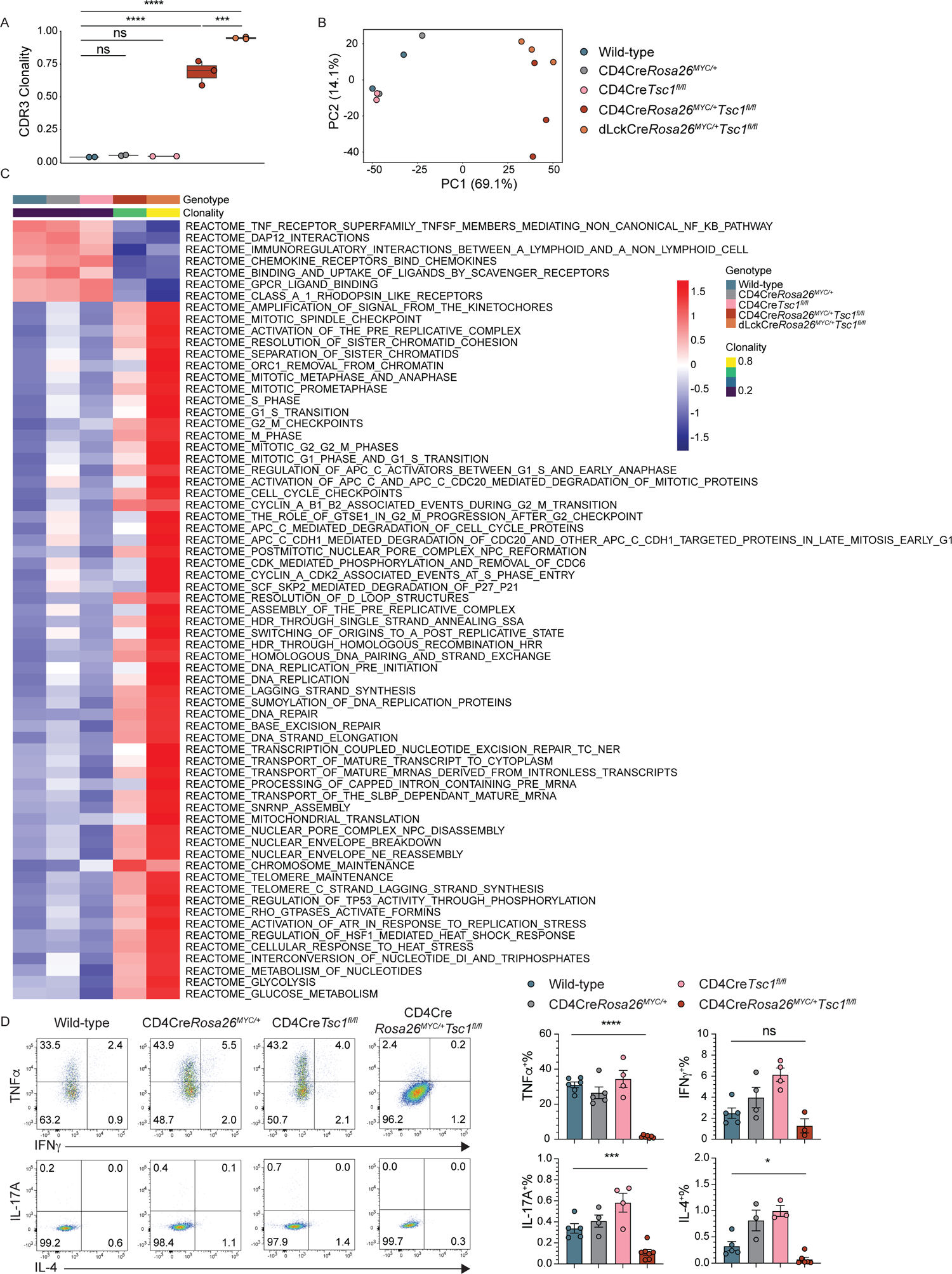
Clonally expanded T cells with constitutive *MYC* transcription and *Tsc1* deletion display decreased effector functions. Related to Figure 2. (A) Box plots of CDR3 clonality for wild-type, CD4Cre*Rosa26^MYC/+^*, CD4Cre*Tsc1^fl/fl^*, and CD4Cre*Rosa26^MYC/+^Tsc1^fl/fl^*, and dLckCre*Rosa26^MYC/+^Tsc1^fl/fl^* mice. (B) Principal Component Analysis (PCA) plot of top two principal components computed from bulk RNA-sequencing of sorted CD4^+^ T cells from wild-type, CD4Cre*Rosa26^MYC/+^*, CD4Cre*Tsc1^fl/fl^*, and CD4Cre*Rosa26^MYC/+^Tsc1^fl/fl^*, and dLckCre*Rosa26^MYC/+^Tsc1^fl/fl^* mice. Percent of variance explained is shown in parentheses for each component. (C) Heatmap of z-scored ssGSEA scores (averaged across 2-3 mice per condition) for select Reactome signature gene sets. Gene sets included are those differentially enriched between wild-type and dLckCre*Rosa26^MYC/+^Tsc1^fl/fl^*T cells, but for heatmap visualization CD4Cre*Rosa26^MYC/+^*, CD4Cre*Tsc1^fl/fl^*, and CD4Cre*Rosa26^MYC/+^Tsc1^fl/fl^*T cells are also shown. Genotypes are ordered horizontally by average CDR3 clonality, and gene sets are manually ordered vertically by functional annotation. (D) Representative flow cytometric analysis (left panel) and frequencies (right panel) of splenic CD4^+^ effector T cells expressing cytokines TNFα, IFNγ, IL-17A, or IL-4 from wild-type, CD4Cre*Rosa26^MYC/+^*, CD4Cre*Tsc1^fl/fl^*, and CD4Cre*Rosa26^MYC/+^Tsc1^fl/fl^* mice after 4 hours PMA/Ionomycin stimulation. All bar graphs are shown as mean ± SEM, each dot representing one mouse. Data are pooled from multiple experiments. [A: one-way ANOVA with Tukey’s multiple comparisons test; D: unpaired t-test, two-tailed, * = p<0.05, ** = p<0.01, *** = p<0.001, **** = p<0.0001, “ns” = not significant.]

**Figure S3.**
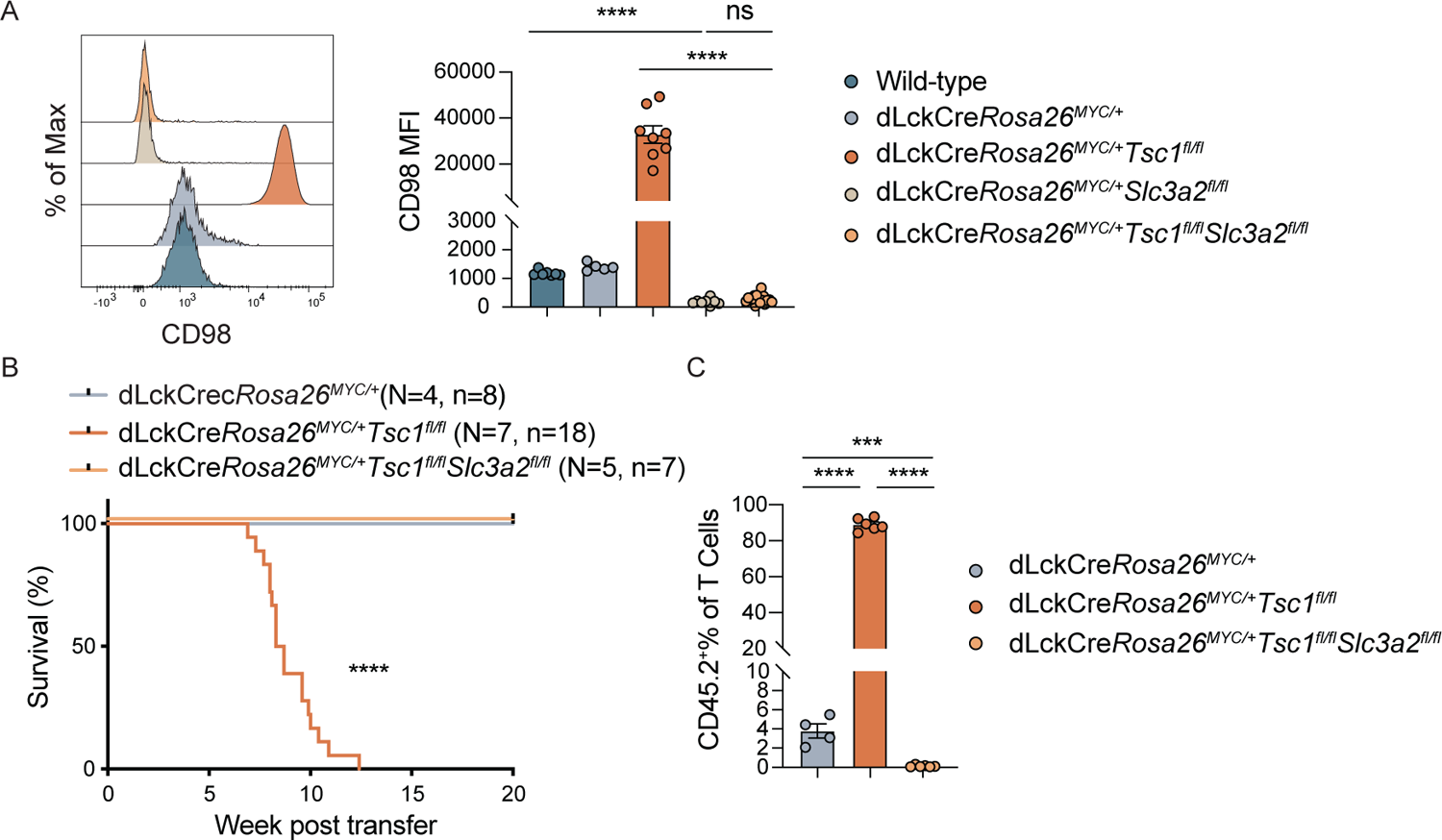
SLC3A2 deficiency corrects the transferrable lethal phenotype of PTCL cells. Related to Figure 3. (A) Representative flow cytometric analysis of CD98 protein staining (left panel) and MFI quantification (right panel) in splenic CD4^+^ T cells from wild-type mice, hCD2^+^CD4^+^ T cells from dLckCre*Rosa26^MYC/+^*, dLckCre*Rosa26^MYC/+^Tsc1^fl/fl^*, dLckCre*Rosa26^MYC/+^Slc3a2^fl/fl^* and dLckCre*Rosa26^MYC/+^Tsc1^fl/fl^Slc3a2^fl/fl^*mice at 13 weeks of age. (B) Survival curves for sub-lethally irradiated recipient mice (CD45.1/CD45.2) after receiving donor (CD45.2) hCD2^+^CD4^+^CD25^-^ T cells from dLckCre*Rosa26^MYC/+^* and dLckCre*Rosa26^MYC/+^Tsc1^fl/fl^*mice, or hCD2^+^CD98^-^CD4^+^CD25^-^ T cells from dLckCre*Rosa26^MYC/+^Tsc1^fl/fl^Slc3a2^fl/fl^*mice. (C) Quantification of dLckCre*Rosa26^MYC/+^*, dLckCre*Rosa26^MYC/+^Tsc1^fl/fl^*, and dLckCre*Rosa26^MYC/+^Tsc1^fl/fl^Slc3a2^fl/fl^* donor T cell frequencies among total splenic TCRβ^+^ T cells determined by the endpoint for recipient mice receiving dLckCre*Rosa26^MYC/+^Tsc1^fl/fl^* donor T cells. All bar graphs are shown as mean ± SEM, each dot representing one mouse. Data are pooled from multiple experiments. [A, C: one-way ANOVA with Tukey’s multiple comparisons test; B: Log-rank (Mantel Cox) test, * = p<0.05, ** = p<0.01, *** = p<0.001, **** = p<0.0001, “ns” = not significant.]

**Figure S4.**
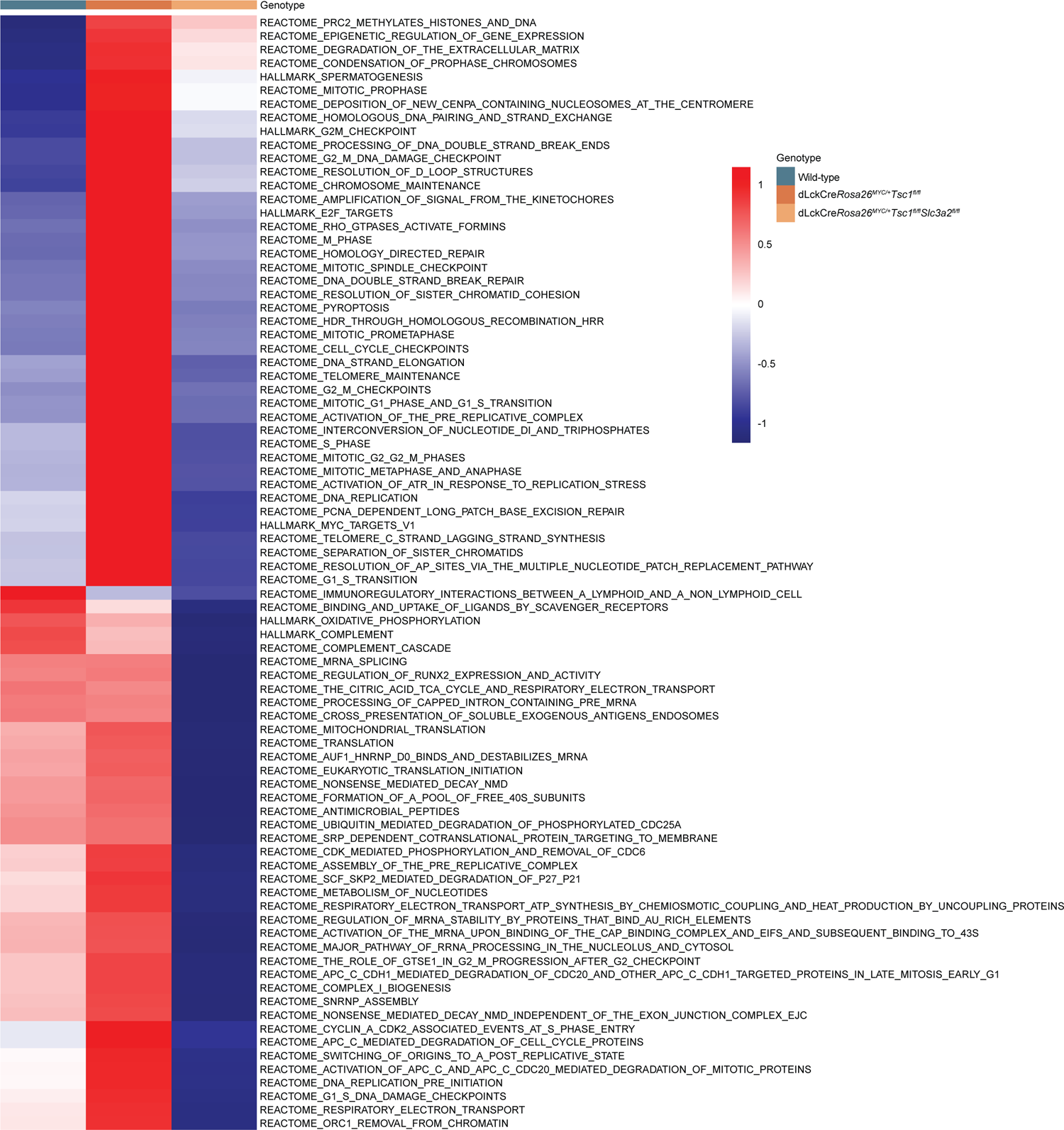
SLC3A2-deficient PTCL cells are transcriptionally distinct. Related to. Figure 4. Heatmap of z-scored ssGSEA scores (averaged across 2-3 mice per condition) for select Hallmark and Reactome signature gene sets. Gene sets included are those differentially enriched between CD4^+^ T cells from 8-week-old dLckCre*Rosa26^MYC/+^Tsc1^fl/fl^*, and dLckCre*Rosa26^MYC/+^Tsc1^fl/fl^Slc3a2^fl/fl^* mice, but for heatmap visualization wild-type mice are also shown. Gene sets are ordered vertically by hierarchical clustering (dendrogram not shown).

**Figure S5.**
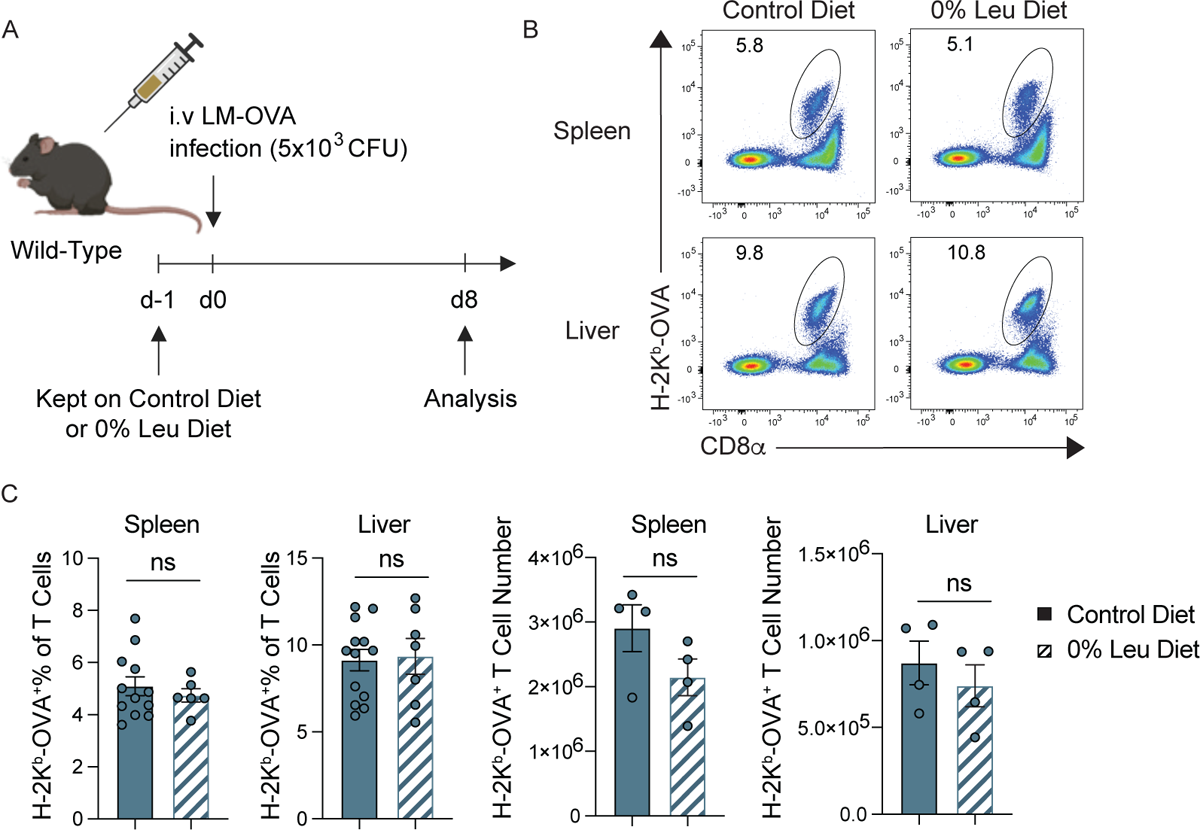
Effector CD8^+^ T cell responses to infection is not impaired by leucine-deficient diet. Related to Figure 5. (A) Schematic illustrating experiments for assessing the impact of a leucine-deficient diet on effector CD8^+^ T cell responses 8 days post LM-OVA infection. Wild-type mice were fed with control diet or 0% leucine diet beginning one day prior to infection with LM-OVA. (B) Representative flow cytometric analysis of H-2K^b^-OVA^+^CD8^+^ T cells in spleen and liver from wild-type mice fed with control diet or 0% leucine diet. (C) Quantification of H-2K^b^-OVA^+^CD8^+^ T cell frequencies among total T cells (left) and numbers (right) in spleen and liver from wild-type mice fed with control diet or 0% leucine diet. All bar graphs are shown as mean ± SEM, each dot representing one mouse. Data are pooled from multiple experiments. (C: unpaired t-test, two-tailed, “ns” = not significant.)

**Figure S6.**
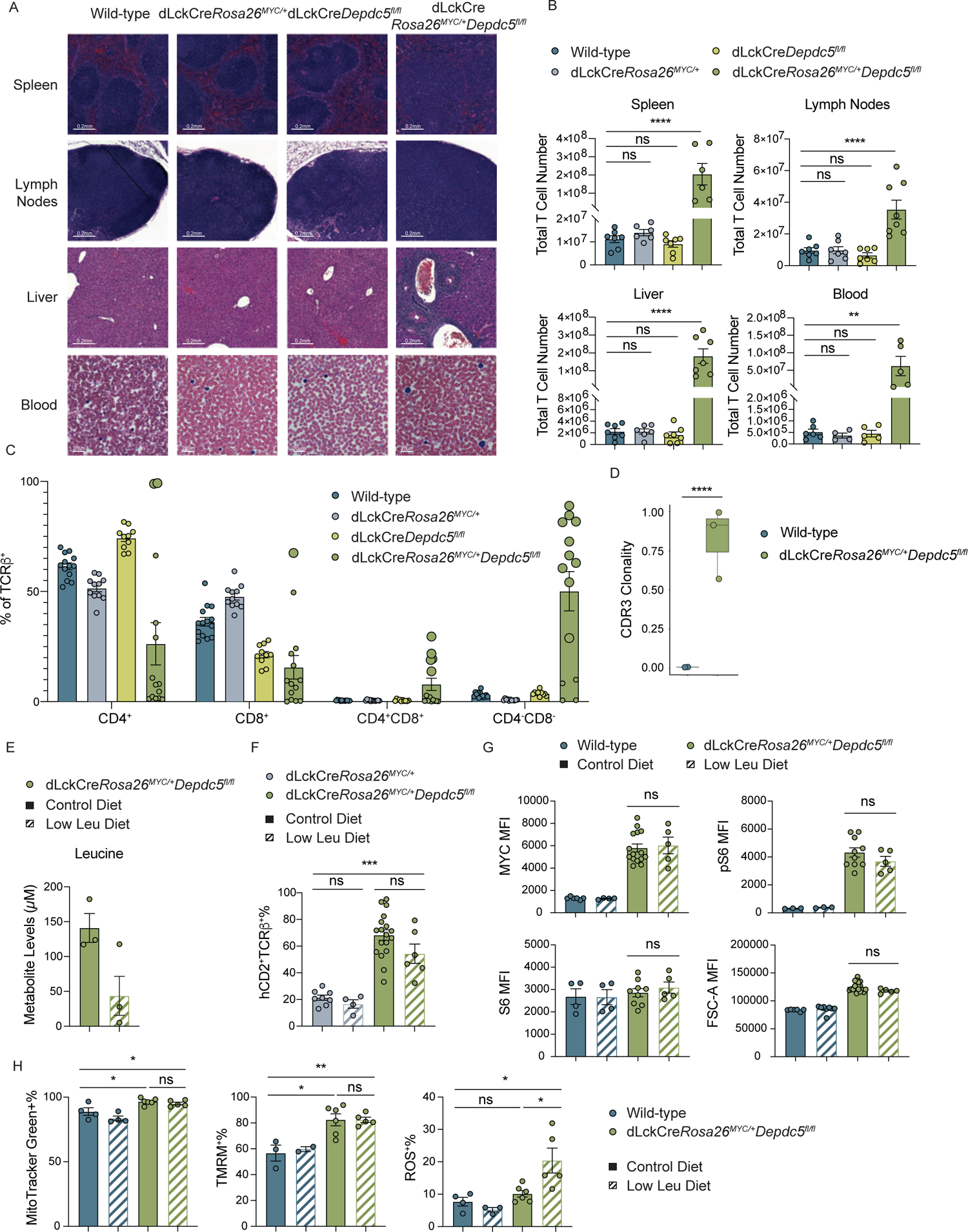
Lethal PTCL driven by constitutive *MYC* transcription and *Depdc5* deletion cannot be rescued by low leucine diet. Related to Figure 6. (A) Rows 1-3: Representative images of hematoxylin and eosin (H&E) staining of spleen, lymph node and liver in wild-type, dLckCre*Rosa26^MYC/+^*, dLckCre*Depdc5^fl/fl^*, and dLckCre*Rosa26^MYC/+^Depdc5^fl/fl^*mice. Original magnification, 21x. Scale bar 0.2mm. Row 4: Representative images of Giemsa stain of blood from wild-type, dLckCre*Rosa26^MYC/+^*, dLckCre*Depdc5^fl/fl^*, and dLckCre*Rosa26^MYC/+^Depdc5^fl/fl^*mice. Original magnification, 63x. Scale bar 20 μm. (B) Quantification of total T cell number in spleen, lymph nodes, liver and blood from wild-type, dLckCre*Rosa26^MYC/+^*, dLckCre*Depdc5^fl/fl^*, and dLckCre*Rosa26^MYC/+^Depdc5^fl/fl^* mice. (C) Frequencies of CD4^+^, CD8^+^, CD4^+^CD8^+^ and CD4^-^CD8^-^ T cells in spleen from wild-type, dLckCre*Rosa26^MYC/+^*, dLckCre*Depdc5^fl/fl^*, and dLckCre*Rosa26^MYC/+^Depdc5^fl/fl^* mice. Enlarged dots represent statistics of expanded T cell populations. (D) Box plots of CD4^+^ T cell CDR3 clonality for wild-type and dLckCre*Rosa26^MYC/+^Depdc5^fl/fl^* mice. (E) Serum leucine levels in dLckCre*Rosa26^MYC/+^Depdc5^fl/fl^* mice fed with control or low leucine diet for 4 weeks. (F) Quantification of splenic hCD2^+^TCRβ^+^ T cell percentage from dLckCre*Rosa26^MYC/+^*and dLckCre*Rosa26^MYC/+^Depdc5^fl/fl^* mice fed with control or low leucine diet for at least 4 weeks. (G) MFI quantification of MYC (upper left panel), phosphorylated S6 (upper right panel), and S6 protein staining (bottom left panel) as well as FSC-A (bottom right panel) in splenic CD4^+^ T cells from wild-type mice and hCD2^+^CD4^+^ T cells from dLckCre*Rosa26^MYC/+^Depdc5^fl/fl^*mice fed with control or low leucine diet for at least 4 weeks. (H) Percentage quantification of MitoTracker Green (left panel), TMRM (middle panel), and MitoSoxRed (right panel) staining in splenic CD4^+^ T cells from wild-type mice and hCD2^+^CD4^+^ T cells from dLckCre*Rosa26^MYC/+^Depdc5^fl/fl^*mice fed with control or low leucine diet for at least 4 weeks. All graphs are shown as mean ± SEM, each dot representing one mouse. Data are pooled from multiple experiments. (B, F, H: one-way ANOVA with Tukey’s multiple comparisons test; D, E, G: unpaired t-test, two-tailed, * = p<0.05, ** = p<0.01, *** = p<0.001, **** = p<0.0001, “ns” = not significant.)

**Figure S7.**
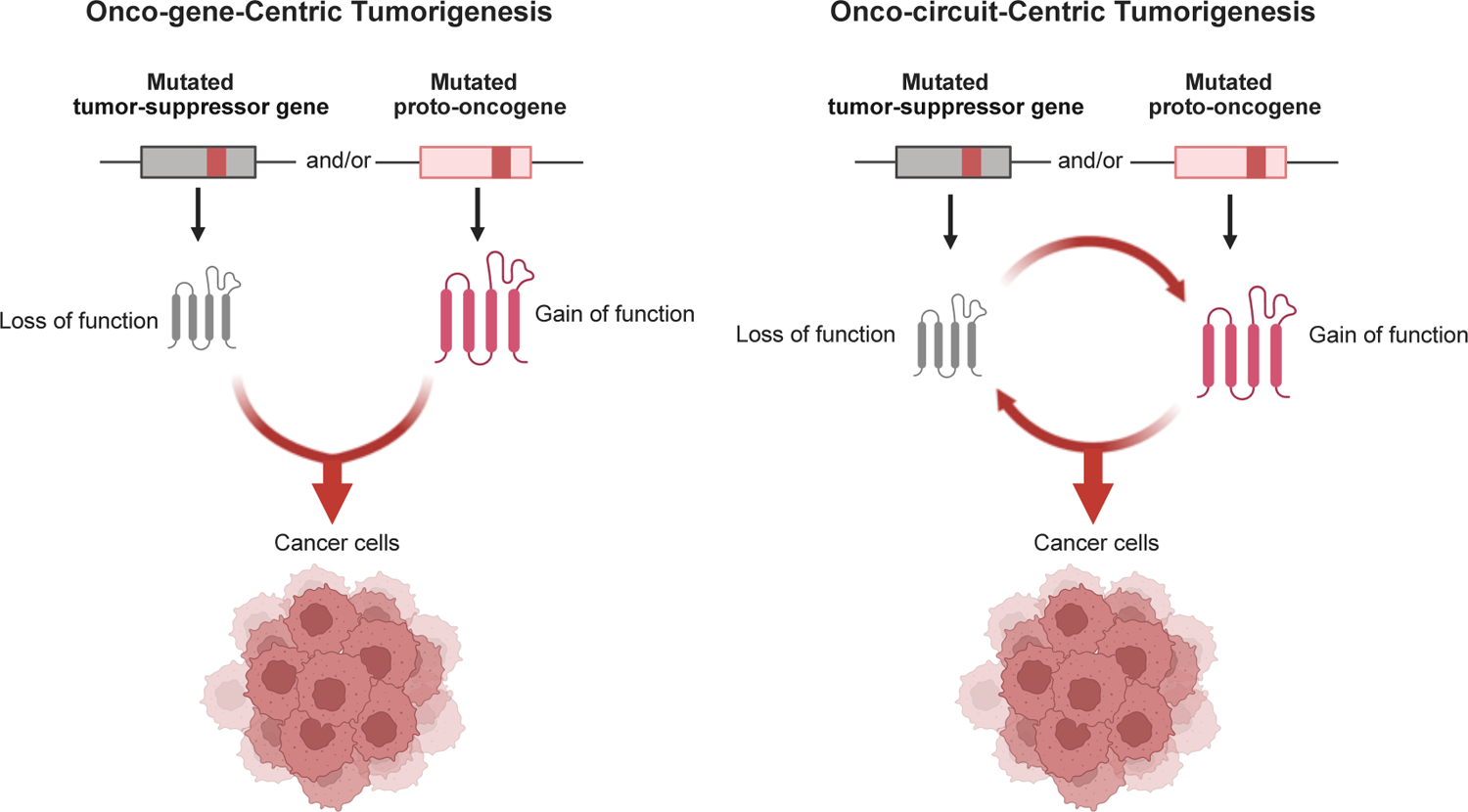
Schematic illustrating onco-gene-centric tumorigenesis and onco-circuit-centric tumorigenesis. Related to Figure 7. The conventional model of onco-gene-centric tumorigenesis proceeds with mutated tumor-suppressor genes and/or mutated proto-oncogenes (i.e. onco-genotype) contributing to cell transformation as independently manifested, but synergistic functional programs (i.e. onco-phenotype). In onco-circuit-centric tumorigenesis, mutated tumor-suppressor genes and/or proto-oncogenes cooperate to enable the manifestation of respective functional programs in a positive feedback loop, which is further engaged to promote cell transformation. Non-mutated cancer cell and environmental components can be integral parts of the onco-circuit, rendering them a new class of cancer therapeutic targets.

Supplemental Table S1: Genes and pathways in T cells differentially expressed between dLckCre*Rosa26^MYC/+^Tsc1^fl/fl^*and wild-type mice.

Supplemental Table S2: Genes and pathways in T cells differentially expressed between dLckCre*Rosa26^MYC/+^Tsc1^fl/fl^*and dLckCre*Rosa26^MYC/+^Tsc1^fl/fl^Slc3a2^fl/fl^*mice.

